# More widespread and rigid neuronal representation of reward expectation underlies impulsive choices

**DOI:** 10.1101/2024.04.11.588637

**Authors:** Rhiannon L. Cowan, Tyler Davis, Bornali Kundu, Shervin Rahimpour, John D. Rolston, Elliot H. Smith

## Abstract

Impulsive choices prioritize smaller, more immediate rewards over larger, delayed, or potentially uncertain rewards. Impulsive choices are a critical aspect of substance use disorders and maladaptive decision-making across the lifespan. Here, we sought to understand the neuronal underpinnings of expected reward and risk estimation on a trial-by-trial basis during impulsive choices. To do so, we acquired electrical recordings from the human brain while participants carried out a risky decision-making task designed to measure choice impulsivity. Behaviorally, we found a reward-accuracy tradeoff, whereby more impulsive choosers were more accurate at the task, opting for a more immediate reward while compromising overall task performance. We then examined how neuronal populations across frontal, temporal, and limbic brain regions parametrically encoded reinforcement learning model variables, namely reward and risk expectation and surprise, across trials. We found more widespread representations of reward value expectation and prediction error in more impulsive choosers, whereas less impulsive choosers preferentially represented risk expectation. A regional analysis of reward and risk encoding highlighted the anterior cingulate cortex for value expectation, the anterior insula for risk expectation and surprise, and distinct regional encoding between impulsivity groups. Beyond describing trial-by-trial population neuronal representations of reward and risk variables, these results suggest impaired inhibitory control and model-free learning underpinnings of impulsive choice. These findings shed light on neural processes underlying reinforced learning and decision-making in uncertain environments and how these processes may function in psychiatric disorders.

## Introduction

Adaptive decision-making in an uncertain and complex world involves predicting and evaluating outcomes and then refining subsequent predictions and actions to improve outcomes. Several theories have formalized this iterative reinforcement learning process as a general learning algorithm^1^ that has inspired much of the super-human performance on unsupervised learning tasks in Artificial Intelligence (AI).^2–5^

The biological neural implementation of reward-based, reinforcement learning is thought to be driven by a dopaminergic network associated with the ventral tegmental area and basal ganglia^6^ within the nucleus accumbens,^7–10^ which project to insula^11,12^ and widely throughout the prefrontal cortex.^13–15^ A reward prediction error (*RPE*) signal was initially observed in primate dopamine neurons.^16^ Dopamine neuron activity is elicited in response to a reward cue, if available, rather than the reward itself, but is suppressed if no reward is presented after a cue (e.g., Pavlovian effect).^17^ Consequently, extensive work has been conducted emphasizing dopamine-mediated RPE as a neural teaching signal to update reward predictions.^7,12,18–21^

Subtle failures in this learning process have been shown to underlie features of maladaptive decision-making and psychiatric disorders.^22,23^ A salient example constitutes one aspect of the multifaceted psychological construct of impulsivity: impulsive choice.^24,25^ Impulsive choice is the tendency to favor smaller, immediate rewards over larger, delayed rewards^26^ and is a central component of bipolar, substance use, anorexia nervosa, and other psychiatric disorders.^22,27^ Stemming from classic reward behavior studies, such as the marshmallow experiment,^28^ an expansive field of research is focused on the neural computations of impulsive choice and its consequences for psychiatric disorders.^29–32^ Numerous studies have highlighted the neural signals correlated with reward expectation in the rodent brain^33,34^ and several research studies extending to reward and impulsive choice research in humans, utilizing intracranial recordings^35–37^ and fMRI.^38^

We extend this work here by studying choice impulsivity with temporal difference (TD) learning models of behavior and brain activity. These models estimate a participants reward expectation and RPE on each trial. Moreover, we study several variants of RPE: signed RPE, unsigned RPE, and asymmetric RPEs. Signed RPE includes the full range of positive and negative RPEs, unsigned RPE is the absolute magnitude of the signed RPE, and asymmetric RPE models positive and negative RPEs separately. Unsigned RPE models unvalenced reward surprise,^39^ whereas asymmetric RPEs can be used to distinguish regions associated with risk sensitivity,^40^ processing of negative outcomes,^41^ and subjective information value around uncertainty.^42^ We show that impulsive choices are characterized by more widespread and inflexible neuronal representations of reward expectation. These results provide insight into the neuronal underpinnings of maladaptive choice behavior underlying numerous psychiatric disorders.

## Results

### Impulsive choice behavior

We recorded neural activity from 43 participants who were undergoing invasive neuromonitoring for surgical treatment of drug-resistant epilepsy (21 female, *M* = 36 ± 10 years of age) while they completed 45 total sessions of BART. The aim of BART is to inflate a simulated balloon, without the balloon popping, to gain points that are linearly related to the size of the balloon, and therefore inflation time (Figure 1a). BART consists of three types of trials: (1) active trials with three color categories of balloon: yellow, orange, and red, which pop at decreasing diameters, (2) passive rewarded trials, where participants do not actively start and stop balloon inflation, but can learn about potential inflation time distributions for each balloon color, and (3) passive unrewarded trials, where the balloons are grey and inflate to a random size and no points are banked. Each subject completed an average of 233.24 (± 23.81) total trials, 81.6 ± 8.32 of which were passive trials. Subjects achieved a mean task performance of 83.38% (± 6.72) and scored an average of 46308 (± 5158) total points comprising of 25820 (± 3728) actively acquired points and 20487 (± 2149) passively acquired points (Figure 1h). There was a statistically significant difference in accuracy among balloon colors (*F*(2, 134) = 75.27, *p* < 10^−21^). Post-hoc tests found that the mean accuracy for balloon color was significantly different between yellow (86.82% ± 8.37) and red balloon accuracies (65.70% ± 13.16), (*p* < 10^−5^, 95% C.I. = 33.74, 72.39) and between red and orange balloon (89.51% ± 7.86) accuracies (*p* < 10^−5^, 95% C.I. = 43.87, 82.52) (see Figure 1g).

**Figure 1.**
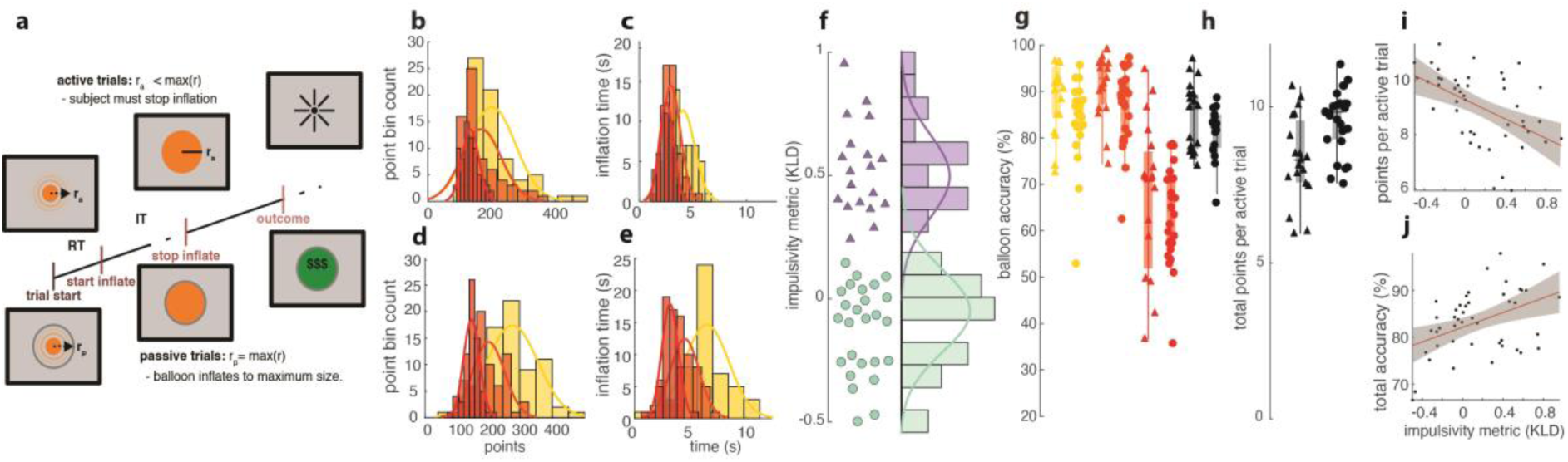
BART behavior reveals a bimodally distributed –reward-accuracy tradeoff underlying impulsive choice. **(a)** Schematic of the BART task. Example trial timelines with an (unsuccessful) active trial shown above and a passive trial shown below. r_a_ and r_p_ indicate active and passive balloon inflation radii, respectively. RT and IT indicate reaction time and inflation time stages, respectively. **(b-c)** Histograms representing the point count distributions (b) and inflation time distribution (c) for red, orange, and yellow balloons for a representative more impulsive (MI) chooser (log Kullback-Liebler Divergence (KLD) score = 0.406), showing similar distributions for all balloon colors. **(d-e)** Histogram representing the point count distributions (d) and inflation time distributions for red, orange, and yellow balloons for a representative less impulsive (LI) chooser (log KLD score = −0.364), showing relatively separate inflation time distributions for all balloon colors. **(f)** A scatter plot of log KLD scores across participants next to histograms. The Gaussian mixture models are overlaid in purple and teal. Triangles represent MI choosers (N = 19); circles represent LI choosers (N = 26). **(g)** Task accuracy (%) for both MI and LI groups for yellow, orange, and red balloons, as well as for total accuracy (black). **(h)** Average total points per active trial for MI and LI subjects (e.g., 10 = 10^3^). **(i)** Regression highlighting significant anticorrelation between impulsivity scores and points during active trials. **(j)** Regression highlighting significant correlation between impulsivity scores and total task accuracy.

There was no significant difference between yellow and orange balloon accuracies (*p* = 0.42). No differences were seen for average active response times (i.e., the time between the cue appearance and the onset of balloon inflation, implemented by the subject), between balloon color categories (*p* = 0.98).

To study impulsive choice, we operationally defined a participant’s propensity to choose a smaller more immediate reward as the distributional distance between active trial inflation time distributions, and the optimal passive trial inflation time distributions (Supplementary Figure S1a & b). We measured inflation time distributional similarity with the Kullback-Leibler Divergence (KLD) between active and passive trial inflation time distributions but found similar results with inflation time distribution means (see Supplementary Figure S3). In examining this measure of IC, we noticed a bimodal distribution of impulsive choosers (see Figure 1f). We therefore fit a gaussian mixture model to impulsivity scores to classify subjects into less-impulsive (LI, *N* = 26, log KLD*_M_* = −0.12) or more-impulsive (MI, *N* = 19, log KLD*_M_* = 0.53) choosers. Our assumption that MI choosers would opt for smaller, more immediate rewards was apparent in the active versus passive trial point and inflation time distributions, and between balloon color point distributions (e.g., Figure 1b-e). Active inflation times were significantly shorter for MI subjects (*M* = 3.68s ± 0.73) than LI subjects (*M* = 4.30s ± 0.50), (*z* = −2.93, *p* = 0.0034). As expected, no differences were seen for passive trial inflation times between groups (MI*_M_* = 6.74s ± 0.34; LI*_M_* = 6.63s ± 0.24; *z* = −1.76, *p* = 0.079).

There was a statistically significant correlation between total points gained during the task and impulsivity scores (adjusted-*R*^2^ = 0.22, *F*(2, 45) = 13.1, *p* = 0.00076). These behavioral results indicate that higher impulsivity scores predicted a reduction in total points throughout the task (see Figure 1i). To control for possible variance in our impulsivity metric over subjects, we also regressed the z-score difference between active and passive inflation time means against KLD scores (adjusted-*R*^2^ = 0.38, *F*(2, 45) = 27.50, *p* = 4.51 × 10^−6^) (Supplementary Figure S3b). Furthermore, the tendency for impulsive choosers to opt for smaller, more immediate reward was also reflected in total point accumulation during the task; indicating that LI subjects gained more active points (*M* = 27270 ± 2782), compared to MI subjects (*M* = 23836 ± 4003), (*z* = −2.86, *p* = 0.0042) (see Figure 1h).

We therefore noticed a score-accuracy tradeoff that subserved the impulsive choice construct in BART performance. Impulsivity scores predicted both points scored in BART (Figure 1i; adjusted-*R*^2^ = 0.24, *F*(2, 45) = 14.9, *p* = 0.00037), with greater impulsivity predicting a reduction in active trial points (*β* = −1790.20), and accuracy (Figure 1j; adjusted-*R*^2^ = 0.17, *F*(2, 45) = 9.83, *p* = 0.0031) with increased impulsivity predicting greater task accuracy (*β* = 7.79). These regression analyses reveal fundamental risk-reward dynamics of BART task behavior such that participants who can resist choosing impulsively exhibit significantly lower mean accuracy yet are more successful overall in number of points awarded (Figure 1i-j & Supplementary Figure S3d).

### Temporal difference modeling of behavior

Having characterized impulsive choosing behavior in BART across trials, we next sought to precisely infer the amount of reward and risk participants predicted on each trial (i.e., value expectation) and how much they were surprised by deviations from that expectation (i.e., prediction error). Therefore, we fit TD learning models to each participant’s behavioral performance on BART and estimated their optimal learning rate parameters (α) for reward (Figure 2) and risk (Figure 3) TD variables using maximum likelihood estimation (see Methods).^18,43^ We updated reward models with actual reward outcomes from the previous trial and risk models were updated by the cumulative probability of success for each type of balloon starting from 0.5 (n.b., for control trials risk = 0). We updated prediction error based on previous trial risk and reward outcomes. Estimating the optimal α for each participant allowed us to infer how learning rate related to impulsivity scores and depended on outcomes. However, we found no significant differences in mean α’s between impulsivity groups for both risk and reward TD models (both *p* > 0.05; Figure 2k & Figure 3k).

**Figure 2.**
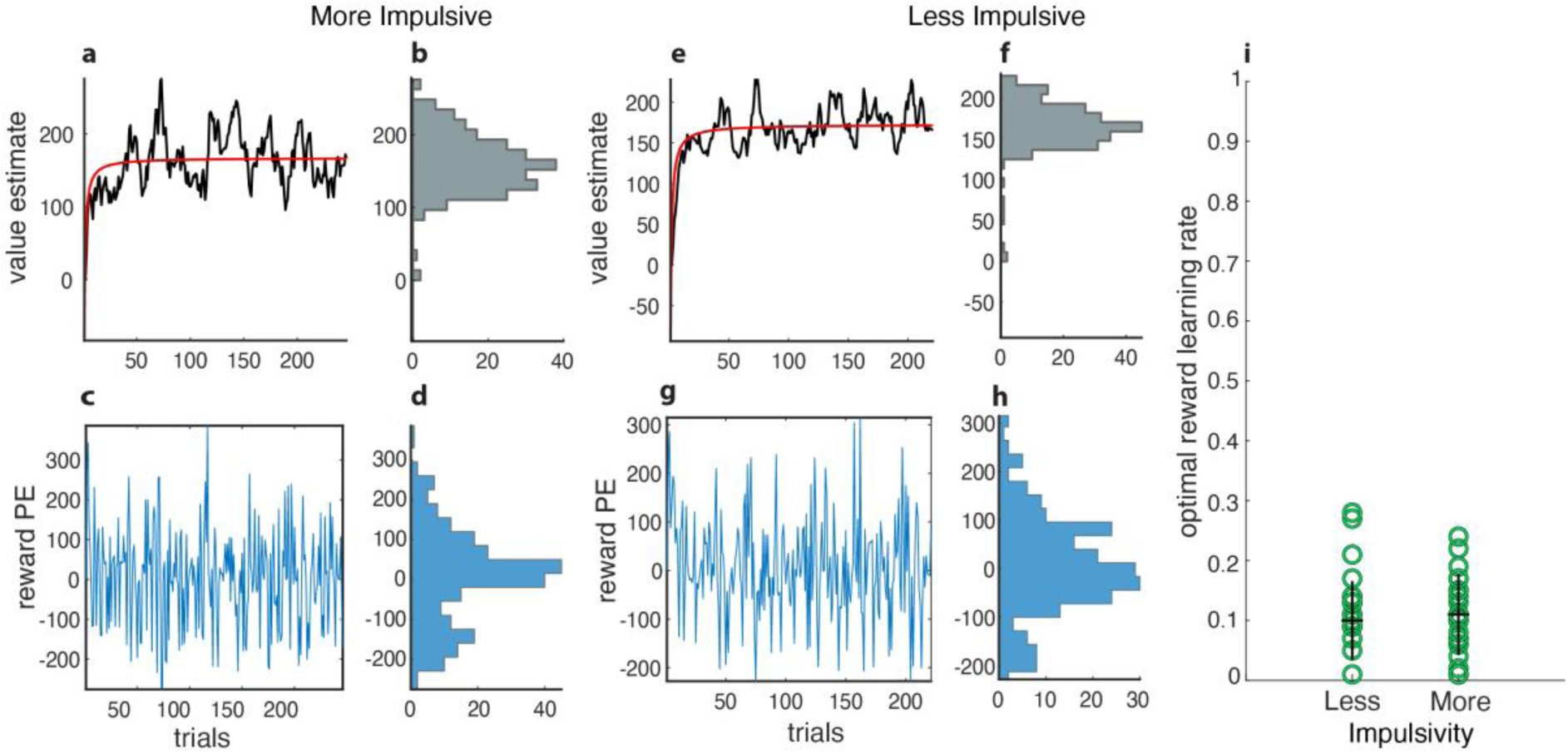
Temporal Difference Learning Models for Reward Variables. **(a-d)** MI subject example. **(a)** Reward Value Expectation (*RVE*) estimates for each trial. **(b)** Histogram count for *RVE* estimates. **(c)** Reward Prediction Error (*RPE*) estimates for each trial. **(d)** Histogram count for *RPE* estimates. **(e-h)** LI subject example. **(e)** *RVE* estimates for each trial. **(f)** Histogram count for *RVE* estimates. **(g)** *RPE* estimates for each trial. **(h)** Histogram count for *RPE* estimates. **(i)** Optimal learning rates for reward variables for MI and LI groups. Each circle represents one participant, and the black crosses represent mean, and standard deviations across participants within each group.

**Figure 3.**
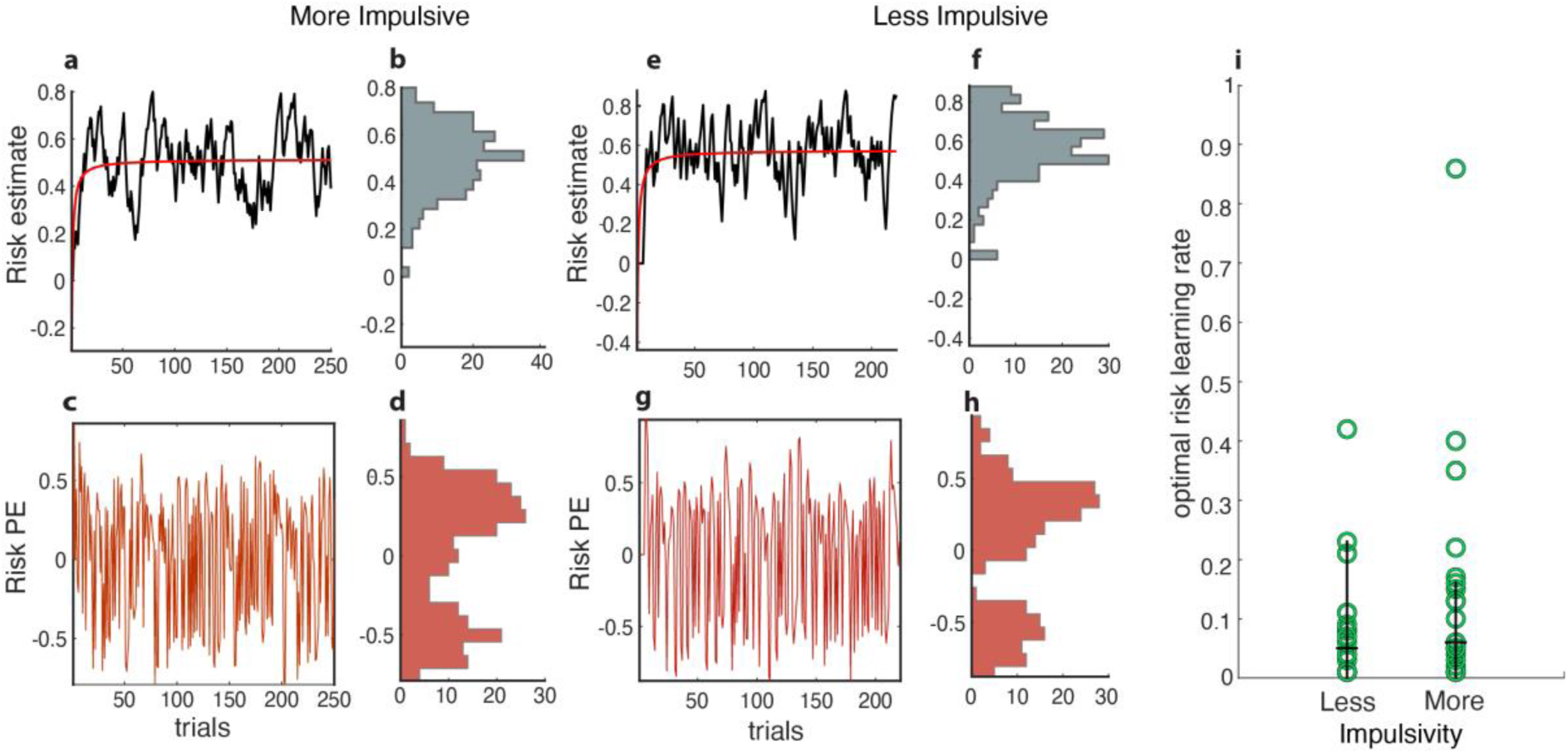
Temporal Difference Learning Models for Risk Variables. **(a-d)** MI subject example. **(a)** *Risk VE* estimates for each trial. **(b)** Histogram count for *Risk VE* estimates. **(c)** *Risk PE* estimates for each trial. **(d)** Histogram count for *Risk PE* estimates. **(e-h)** LI subject example. **(e)** *Risk VE* estimates for each trial. **(f)** Histogram count for *Risk VE* estimates. **(g)** *Risk PE* estimates for each trial. **(h)** Histogram count for *Risk PE* estimates. **(i)** Optimal learning rates for risk variables for MI and LI groups. Each circle represents a patient and black crosses represent means and standard deviations.

### Greater neural encoding of reward than risk when modelling temporal difference learning variables

We observed a significant behavioral relationship between BART task performance (accuracy and points) and impulsivity level, however there were no significant differences in optimal learning rates estimated for each TD model. This led us to examine the neural correlates of reward and risk expectation and surprise to understand which human brain areas compute reward and risk expectation and surprise during impulsive choices. To do so, we examined broadband high frequency local field potentials (HFA: 70 – 150 Hz) recorded from 3259 stereo-electroencephalography (sEEG) and electrocorticography (ECoG) contacts (*M* = 72.42 ± 17.17 electrodes per participant; 1483 electrode contacts in the MI group and 1359 electrode contacts in the LI group). HFA is an established correlate of aggregate neuronal firing near each electrode.^44–46^

We modeled HFA from each electrode as linear functions of the TD model variables across trials for both value expectation and surprise. We assumed participants estimated the risk associated with a given balloon from its color and therefore used balloon-aligned HFA (time window: 0.25ms – 1.25ms) for risk models. For analogous reasons, we used outcome-aligned HFA (time window: 0.25ms – 1.25ms) for reward models. We therefore modeled neural encoding across trials for reward value expectation (*RVE*), *RPE*, risk value expectation (*Risk VE*) and, risk prediction error (*Risk PE*), while controlling for any HFA responses from the sensory salience of the balloon popping via linear mixed effect models (see Methods). Coefficient models for each variable were used to calculate the significant contacts from the total contacts for each region of interest.

To test whether reward and risk representations were observed in each brain region at levels greater than those expected by chance (5% threshold), a series of proportion tests were conducted. Our proportion analysis accounts for uneven contact numbers in each hemisphere. We had too many electrode contacts to sufficiently represent the contact locations on a 3D brain. To fully visualize the data, we simplified the anatomical representation using dimensionality reduction, showing the contacts on a 2D brain (see Figures 4e & 5e). We found an overall higher proportion of ECoG contacts encoding reward model variables (*RPE* and *RVE* = 79.48%), rather than risk model variables (*Risk PE* and *Risk VE* = 23.45%; *χ*^2^(1) = 1014.42, *p* < 10^−5^). A subsequent linear mixed effects model that controlled for model variance derived from the sensory salience of the balloon pop also found a higher proportion of contacts encoding reward model variables (39.24%), rather than risk model variables (19.74%; *χ*^2^(1) = 150.50, *p* < 10^−5^). In this model, significantly more contacts encoded *RPE* (21.14%) than *RVE* (13.58%; *χ*^2^(1) = 31.68, *p* < 10^−5^). We report results from the salience-controlled model below and the original model results are reported in the supplementary information. Additionally, we examined encoding of asymmetric and unsigned *PEs* to further understand more specific encoding of a variety of *PE* responses. These models included random effects to control for salience of negative outcomes. These results indicate that across all participants, even during a risky decision-making task, many more contacts encoded reward, versus risk, TD variables. In the following two sections we describe how brain networks encode these variables in the context of more impulsive or less impulsive choices.

**Figure 4.**
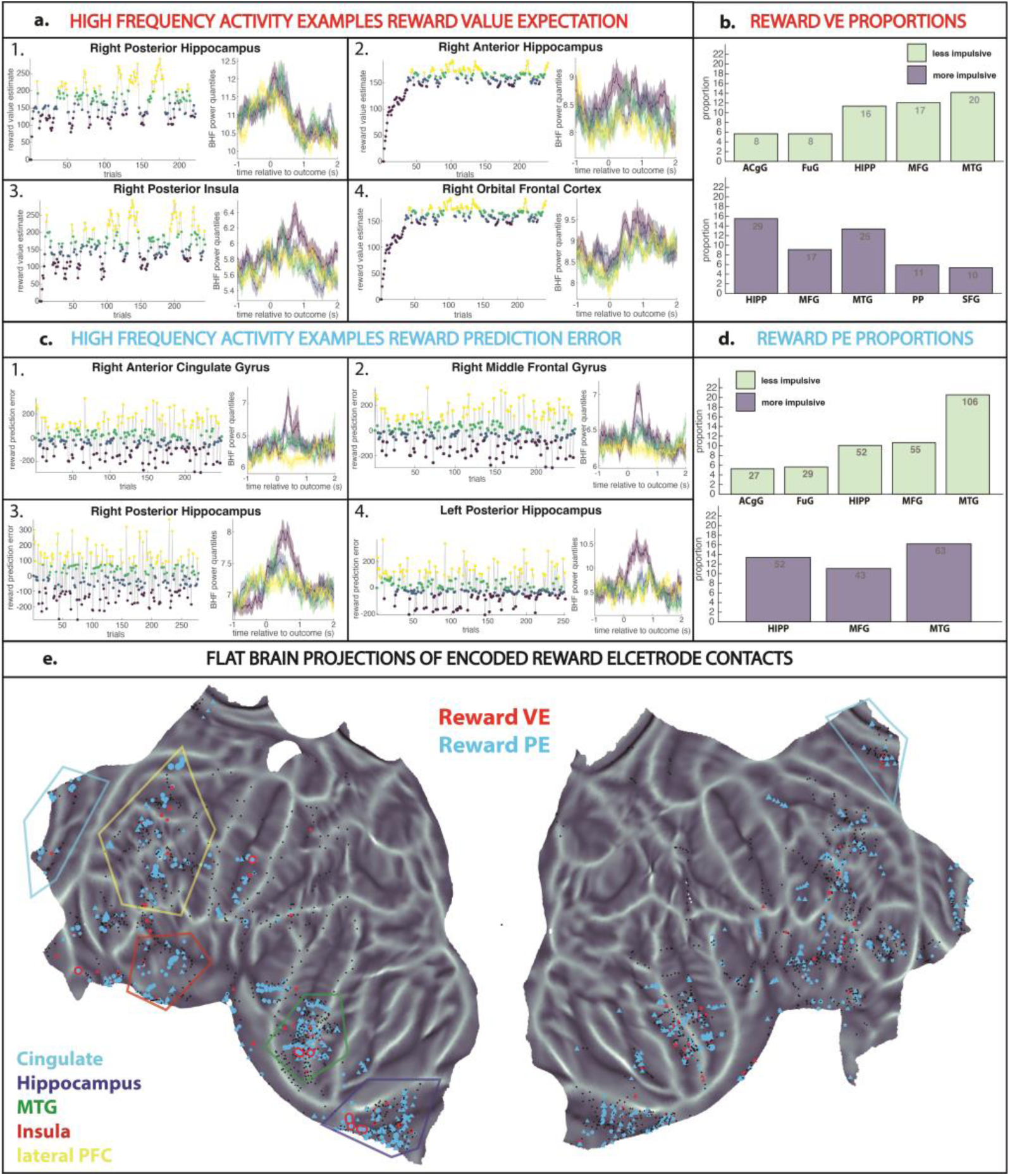
High Frequency Activity and Regional Brain Responses for Reward Variables utilizing the Balloon Pop Control Model. **(a)** Examples of high frequency activity (70-150 Hz) for RVE. Each numbered subplot shows the trajectory of value estimates across trials for a single session, with each trial color-coded by its VE quartile, on the right, and average HFA responses in each quartile, color-coded accordingly, on the left. (1) Right Amygdala (2) Right Ventral Cingulate (3) Right Orbital Frontal Cortex (4) Right Anterior Hippocampus. **(b)** Bar graphs highlighting the proportions of significantly encoded electrode contacts for *RVE* between LI (light green) and MI (light purple) groups. **(c)** Examples of high frequency activity (70-150 Hz) for RPE, organized as in **a**. (1) Right Ventral Cingulate (2) Left Posterior Hippocampus (3) Right Anterior Hippocampus (4) Right Amygdala **(d)** Bar graphs highlighting the proportions of significantly encoded electrode contacts for *RPE* between LI (light green) and MI (light purple) groups. **(e)** Flat brain representation of *RVE* (red) and *RPE* (blue) encoding in the brain.

### Impulsive choosers exhibit greater neural encoding of Reward PE and Reward VE

Overall, for our reward models we saw significantly more contacts that encoded *RPE* (21.14%) than *RVE* (13.58%; *χ*^2^(1) = 31.68, *p* < 10^−5^). In testing for differences in neural encoding of TD reward model variables between impulsivity groups, we found significantly more electrode contacts that encoded *RVE* in MI choosers (8.49%) compared to LI choosers (4.19%; *χ*^2^(1) = 25.84, *p* < 10^−5^). Significantly more contacts also encoded *RPE* in MI choosers (13.71%) compared to LI choosers (9.84%; *χ*^2^(1) = 11.08, *p* < 10^−3^; Figure 4). Hemispheric differences in encoding were similar in this model, showing significantly increased reward encoding in the left hemisphere (22.40%) than in the right hemisphere (13.87%; *χ*^2^(1) = 40.33, *p* < 10^−5^). Unless stated otherwise, all models had significantly more electrode contacts encoding variables in the left hemisphere. When examining the intersection between hemisphere and impulsivity, MI choosers had significantly more reward that encoded contacts in the left hemisphere (15.34%) than the right hemisphere (5.65%; *χ*^2^(1) = 84.13, *p* < 10^−5^), whereas LI choosers had significantly more contacts that encoded reward in the right hemisphere (8.33%) than the left hemisphere (4.48%; *χ*^2^(1) = 19.47, *p* = 10^−5^). These results show that reward expectation and surprise are more widespread in MI choosers specifically.

For the asymmetric TD model, significantly more contacts encoded *negative RPE* (17.17%) than *positive RPE* (13.65%; *χ*^2^(1) = 7.63, *p* < 0.0057). Between impulsivity groups, we found significantly more electrode contacts that encoded *positive RPE* for LI choosers (6.99%) compared to MI choosers (5.16%; *χ*^2^(1) = 4.68, *p* = 0.03), but no significant difference in *negative RPE* encoding was observed between impulsivity groups (MI: 10.32%, LI: 8.67%; *χ*^2^(1) = 2.58, *p* = 0.11). Similarly, we found no difference between the proportion of contacts that encoded unsigned *RPE* for MI choosers (8.15%) compared to LI choosers (5.37%, *χ*^2^(1) = 10.07, *p* = 0.11). These results suggest that LI choosers are more susceptible to *positive RPE* encoding, which may reflect a predisposition for adaptive value estimation.

For both impulsivity groups, *RVE* was primarily encoded in frontotemporal regions: middle temporal gyrus (MTG; 9%), middle frontal gyrus (MFG; 10%), hippocampus (HIPP; 14%), and anterior cingulate gyrus (ACgG; 11%) (Supplementary Figure S6a, S6b, & Table S3). MI choosers additionally encoded *RVE* in Planum Polare (PP; 31%) and superior frontal gyrus (SFG; 29%) and LI choosers additionally encoded *RVE* in fusiform gyrus (FuG; 9%) and ACgG (9%). For both groups, *RPE* was primarily encoded in MTG (33%), MFG (29%), and HIPP (33%). LI choosers additionally encoded *RPE* in FuG (33%) and ACgG (32%) (Figure 4d). When *RVE* and *RPE* were analyzed in combination, common regions that encoded reward included MTG (35%), MFG (30%), HIPP (37%), and ACgG (31%), with the LI group also significantly encoding reward in the entorhinal area (ENT; 40%).

Significant neural encoding of asymmetric model variables had a fairly distinct distribution. *Positive RPE* and *negative RPE* were primarily encoded in frontotemporal regions and HIPP with unique encoding of *positive RPE*s in Amygdala (AMY) (28%) and *negative RPE*s in medial orbital gyrus (MOrG) (30%). Representations of *positive RPE* in MI choosers were restricted to the temporal lobe, in AMY (37%), HIPP (12%), and PP (23%), whereas LI choosers encoded *positive RPE* in AMY (21%), Anterior Orbital Gyrus (AOrG) (32%), ACgG (9%), and ENT (10%).

MI choosers encoded *negative RPE* in MTG (29%), HIPP (38%), MFG (23%), ACgG (20%), and MOrG (42%), whereas LI choosers encoded *negative RPE* in MTG (13%), MFG (22%), HIPP (19%), and ENT (21%). Interestingly, hemispheric encoding revealed that MI choosers had more widespread encoding of *positive RPE* (*χ*^2^(1) = 0.90, *p* = 0.34) which we did not see for LI choosers.

For both groups, significant neural encoding of unsigned *RPE*s was observed in MTG (33%), HIPP (33%), and MFG (29%). LI choosers additionally encoded unsigned *RPE* in FuG (33%) and ACgG (32%) (see Supplementary Materials for all asymmetric and unsigned model results). These results detail hemispheric and regional patterns in the neural representations of reward expectation and surprise across varying levels of impulsive choice, with TD model variables being represented in a frontotemporal network encompassing hippocampus, medial prefrontal, orbitofrontal, and dorsolateral prefrontal cortices.

### Less impulsive choosers encode risk prediction error over risk expectation

The neural computations of risk and uncertainty are likely both interwoven and independently associated with reward encoding in the human brain. Therefore, we next examined the neural encoding of TD model variables related to risk. These models followed the standard temporal difference learning format but were updated by the level of uncertainty around outcome for balloons of the same color (see Methods). Overall, significantly more contacts encoded *Risk PE* (11.24%%) than *Risk VE* (10.32%; *χ*^2^(1) = 0.71, *p* = 0.40). Between impulsivity groups, LI choosers had a greater proportion of contacts encoding *Risk PE* (LI = 7.16%, MI = 4.96%; *χ*^2^(1) = 6.78, *p* = 0.0092) despite observing no differences for *Risk VE* (LI = 4.87%, MI = 4.41%; *χ*^2^(1) = 0.37, *p* = 0.54). Overall, risk was not preferentially encoded in either hemisphere (left hemisphere = 10.93%, right hemisphere 10.74%; *χ*^2^(1) = 0.03, *p* = 0.86). Specifically, *Risk VE* was not significantly encoded to a greater extent in the left hemisphere compared to the right hemisphere for LI choosers (*χ*^2^(1) = 1.69, *p* = 0.19) or MI choosers (*χ*^2^(1) = 0.34, *p* = 0.56). MI subjects had more risk encoding contacts in the left hemisphere (6.45%) than the right hemisphere (2.40%; *χ*^2^(1) = 32.60, *p* < 10^−5^), whereas LI subjects had significantly more risk encoding contacts in the right hemisphere (8.33%) than the left hemisphere (4.48%; *χ*^2^(1) = 19.47, *p* < 10^−5^). No differences were observed in the asymmetric risk model (Supplementary Figure S6c). Overall, these results show that risk surprise encoding is more pertinent and widespread in MI choosers.

For both impulsivity groups, *Risk VE* was encoded in MTG (4%), MFG (10%), HIPP (8%), ACgG (9%), FuG (11%), and MOrG (12%). LI choosers additionally encoded *Risk VE* in ENT (12%) while MI choosers additionally encoded *Risk VE* in Ains (7%) (Figure 5b). Proportion tests revealed *Risk PE* was encoded in MTG (6%), MFG (13%), HIPP (8%), and ACgG (14%). Between impulsivity groups, *Risk PE* was commonly encoded in MTG (6%), MFG (13%), HIPP (8%), and ACgG (15%) with LI choosers and MI choosers additionally encoding *Risk PE* in MOrG (15%) and superior frontal gyrus (SFG; 11%), respectively (Figure 5d). When analyzed in combination, risk variables were commonly encoded in MTG (10%), MFG (19%), HIPP (14%), and ACgG (20%), while MI choosers also encoded risk in Ains (12%) and LI choosers additionally encoded risk in FuG (18%) and MOrG (32%).

**Figure 5.**
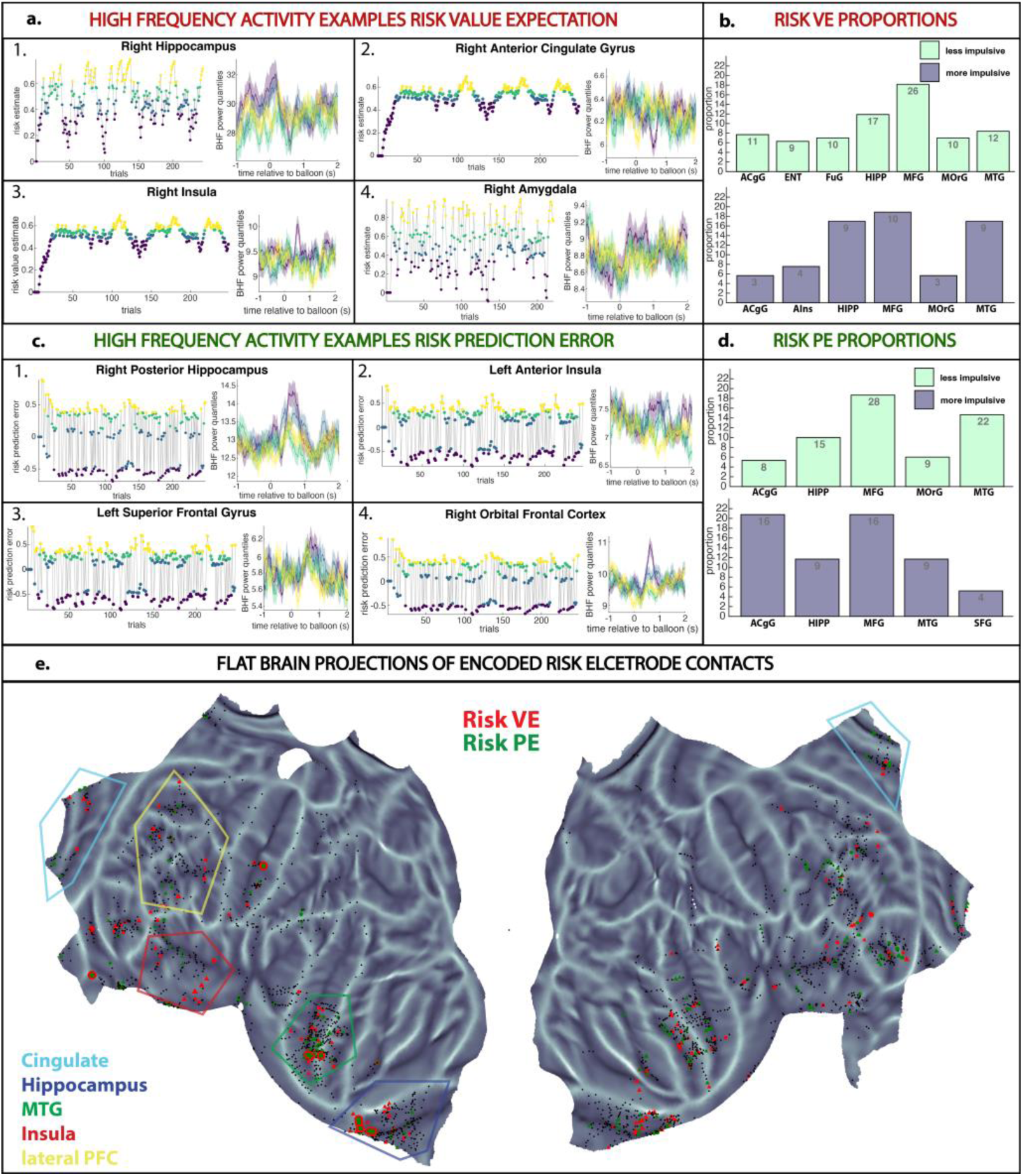
High Frequency Activity and Regional Brain Responses for Risk Variables utilizing the Balloon Pop Control Model. **(a)** Examples of high frequency activity (70-150 Hz) for *Risk VE*. Value estimates across trials highlights quantile expression differences for the fourth quantiles over the first three quantiles. (1) Right Anterior Hippocampus (2) Right Hippocampus (3) Right Amygdala (4) Right Putamen. **(b)** Bar graphs highlighting the proportions of significantly encoded electrode contacts for *Risk VE* between LI (light green) and MI (light purple) groups. **(c)** Examples of high frequency activity (70-150 Hz) for *Risk PE*. Prediction error estimates across trials highlights quantile expression differences for the fourth quantiles over the first three quantiles. (1) Right Insula (2) Right Posterior Hippocampus (3) Right Orbital Frontal Cortex (4) Left Caudate **(d)** Bar graphs highlighting the proportions of significantly encoded electrode contacts for *Risk PE* between LI (light green) and MI (light purple) groups. **(e)** Flat brain representation of *Risk VE* (red) *and Risk PE* (blue) encoding in the brain.

For the asymmetric *Risk PE* model, *positive Risk PE* were encoded in frontotemporal regions, in HIPP (4%), ACgG (10%), ENT (12%), FUG (8%), and MOrG (12%) and *negative Risk PE*s were predominantly encoded in frontotemporal regions, in HIPP (6%), ACgG (7%), MOrG (15%), and Posterior Orbital Gyrus (POrG) (21%). Between impulsivity groups, MI choosers encoded *positive Risk PE* in ACgG (16%), ENT (12%), Ains (10%), Orbital Inferior Frontal Gyrus (OrIFG) (24%), Triangular part of IFG (13%), and PorG (14%), whereas LI choosers encoded *positive Risk PE* in ENT (12%) and MOrG (17%). MI choosers encoded *negative Risk PE* in ACgG (11%), Ains (10%), PP (14%), STG (21%), and POrG (18%), whereas LI choosers encoded in *negative Risk PE* in MOrG (22%), Central Operculus (CO) (22%), and Caudate (23%). Unsigned *Risk PE* was encoded in MTG (6%), MFG (135), HIPP (8%), and ACgG (15%), with LI choosers additionally encoding in MOrG (15%) and MI choosers additionally encoding in SFG (11%) (see Supplementary Figures S7a-c, & Table S4). These results show anatomical differences in neural encoding of Risk expectation and surprise across trials, with LI choosers preferentially encoding Risk related surprise in a frontotemporal neural network including the anterior temporal lobes, ACC, and OFC.

### Post-error response times support more rigid VEs in more impulsive choosers

In the results above, we show that while there is no difference in reward or risk related learning rates between impulsivity groups, there are substantial differences in the neural representations underlying impulsive behavior, with more widespread representations of reward expectation and surprise in MI choosers. We reasoned that these neural representations might manifest themselves in altered behavioral updating across trials. We therefore sought to test the hypothesis that there were significant outcome-dependent changes in response time on subsequent trials. Overall, there was no significant difference in response times between impulsivity groups for active trials (*z =* - 0.91, *p* = 0.36), banked trials (*z =* −1.02, *p* = 0.31), or popped trials (*z =* - 1.02, *p* = 0.41). However, there was a significant slowing of response times in trials after an unrewarded outcome (i.e., popped), for MI choosers compared to LI choosers, shown via a change in post-outcome response times (Δresponse time: MI = 0.061, LI = −0.072; *z* = 2.63, *p* = 0.0085). We interpret this post-pop response time slowing as evidence of a proclivity for MI choosers to have delayed disengagement after unrewarded trials.

We then examined how these behavioral tendencies aligned with reward and risk TD model variables. We opted to test for response time differences relative to two TD model variables: cue-aligned value expectation on the current trial and prediction error from the previous trial. We thus sought to understand how trials with similar value expectations motivate and modulate response time behavior based on prediction error signals. For this analysis we only analyzed trials with similar *RVEs* (median ± 1 SD). To examine if the cue-aligned *VE*s predicted response times between impulsivity groups, we regressed the correlation coefficient of an ANOVA model *VE* and response times amongst balloon colors against impulsivity level and found a significant correlation with MI choosers less likely (*β* = −0.85) to change their response time in response to different reward probability cues (i.e., balloon colors; Figure 6g ; adjusted-*R*^2^ = 0.069, *F*(2, 45) = 4.27, *p* = 0.045). As an additional control, we regressed the *Risk PE* and response time *t*-value on the previous trial against impulsivity level and found a significant correlation with MI choosers more likely to have a negative relationship (*β* = −1.12) between previous trial risk prediction and current trial response time (Figure 6h; adjusted-*R*^2^ = 0.10, *F*(2, 45) = 6.11, *p* = 0.018) (see Supplementary Figures S8 & S9). These results show that for relatively similar reward expectations, general outcome-dependent changes in response time occur, and those changes are parametrically modulated by the reward predictive cue category (i.e. the color of the balloon the participant is about to inflate). Moreover, the magnitude of these changes negatively correlates with impulsivity level across the study cohort, indicating that LI choosers are more likely to implement inhibitory control in response to the reward predictive cue, that is the balloon color.

**Figure 6.**
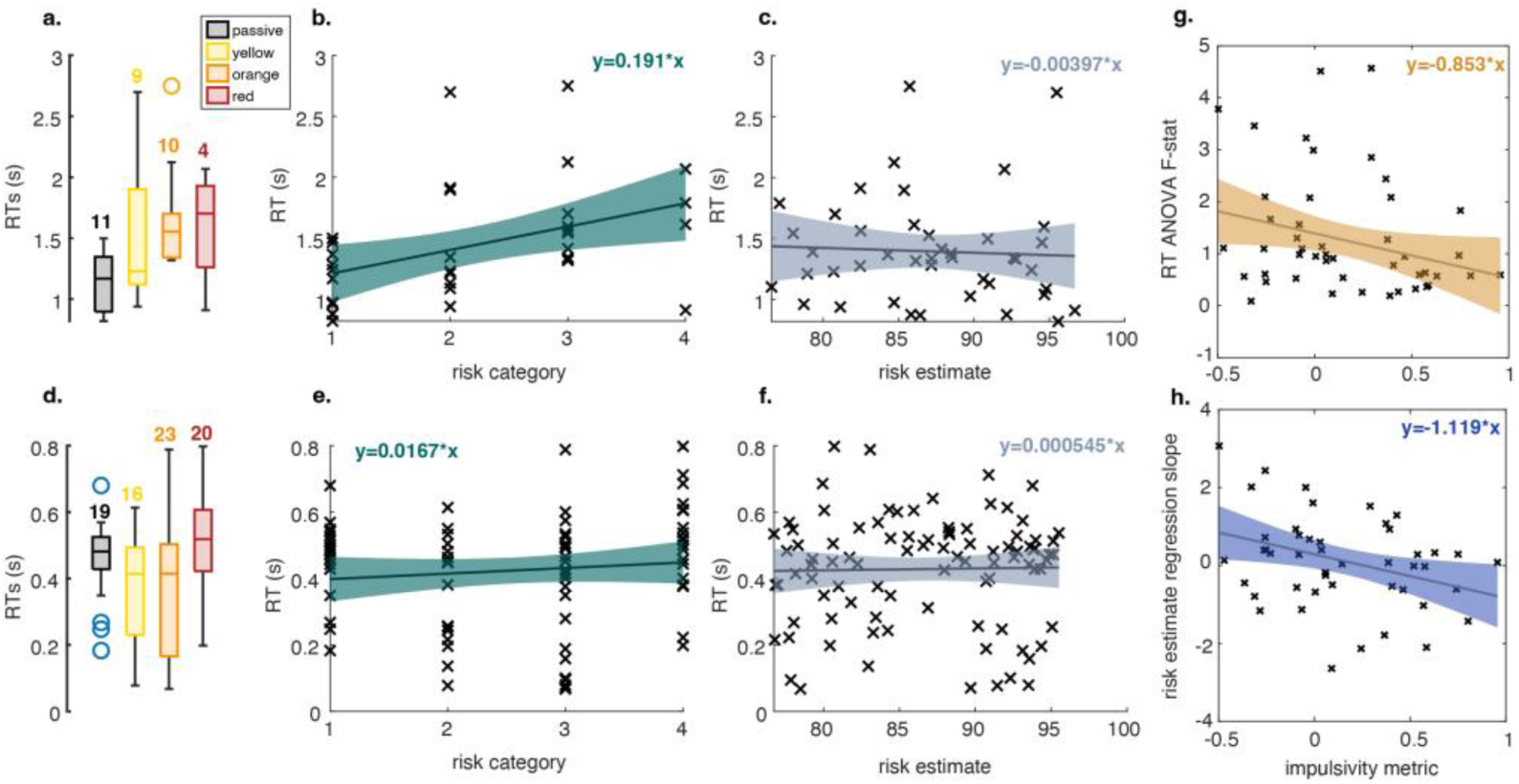
Response Time, Value Expectation, and Prediction Error Interactions. **a-c.** MI example **(a)** Response Times for balloons categories (*F*(3) = 2.8, *p* = 0.05). **(b)** Regression between response times and risk categories (passive, yellow, orange, red). **(c)** Regression between response times and risk value estimate. **d-f.** LI example **(d)** Response Times for balloons categories (*F*(3) = 3.78, *p* = 0.01). **(e)** Regression between response times and risk categories (passive, yellow, orange, red). **(f)** Regression between response times and risk value estimate given. **g-h.** Across-Subjects Value Expectation. **(g)** Calculated *F*-statistic taken from RT VE ANOVA regressed against impulsivity metric. **(h)** Previous trial risk estimate *t*-statistic regressed against impulsivity.

## Discussion

Reward and risk prediction are critical components of naturalistic reinforcement learning and value-based decision-making. However, it remains unknown how reward and risk are encoded in the human brain across time and those processes correspond with impulsive behavior. Previous research has used brain imaging and neuroimaging techniques such as fMRI and scalp EEG, to look at how impulsivity is modulated by reward-related stimuli.^47–49^ We leveraged the spatiotemporal specificity of intracranial recordings from humans to further dissect these processes by exploring how an analog of population neuronal firing represented TD learning algorithms, with specific regard for the implications of impulsive behavior on decision-making. We observed no significant differences in optimal learning rates between impulsivity groups, suggesting no clear differences between how quickly MI and LI choosers learn from deviations in expected reward and risk. However, our findings support the current literature on value-based decision-making and the maladaptive processes associated with impulsive choosing.^50,51^ We generally showed that reward is encoded more than risk, *PE* more than *VE*, and that more impulsive choices are characterized by more widespread and rigid *VE*s.

### Impulsive Choosing

Choice impulsivity is likely linked to an inability to assert inhibitory control over certain actions, thus exacerbating a proclivity for relapse in substance use disorder.^24,52–54^ In this study, we examined how choice impulsivity modulated reward and risk in a cognitive paradigm. As hypothesized, in the behavioral analyses, we observed a trade-off between potential reward and performance accuracy. MI choosers opted for smaller, more immediate rewards (e.g., stopping balloon inflation earlier) and LI choosers opted for riskier, larger rewards but overall gaining more points (i.e., letting the balloon inflate to a larger size to gain more points, increasing the probability of the balloon popping). This reward-seeking behavior coupled with loss aversion results in MI choosers exhibiting suboptimal task performance: fewer points gained relative to the potential gains within the task. On the other hand, less impulsive choosers were able to *optimize* the rewarded outcomes (e.g., high total points gained) and still maintain relatively high accuracy. This behavior displayed by MI choosers may be due to a disinhibitory tendency: once a reward is present, it becomes harder to ‘wait’ for a potentially greater, future reward. This increased sensitivity or propensity to immediate reward, or temporal discounting) is reflective of impulsivity across a number of addictive disorders.^53,55–57^

An alternative explanation is loss aversion,^58^ where impulsive choosers will opt for any reward, even if small, over the risk of losing that reward. Loss aversion is potentially apparent in MI choosers, with tendencies to *maximize* the number of rewarded outcomes (e.g., high task accuracy), independent of the potential reward on a given trial. Supporting this, we found that MI choosers exhibited greater response times on the current trial if the previous trial was unrewarded (i.e., popped balloon). Therefore, MI choosers are more likely to have a negative relationship between previous trial risk prediction. We initially interpreted this as an increased allocation of cognitive effort, post-error, to minimize loss and increase accuracy on prospective trials. Prior findings suggest that increased motor responses (i.e., faster response time) could be related to high value and salience.^59,60^ Despite a correlation between *Risk PE*s on previous loss trials and response times on the current trial, we did not observe adaptive modulation of inflation time on the next trial. Inflation time modulation would suggest that MI choosers are adapting decision-making behaviors to avoid loss, instead they continue with rigid value estimates and low-risk, reward seeking. This behavioral manifestation of rigidity of reward value aligns with the idea that substance-dependent individuals do not adjust behavior after a negative outcome, such as an unrewarded trial.^57^ Recently, a study found that this may reflect abnormal neural processing of negative outcomes, shown via differences in low-frequency (delta-theta) activity relative to losses, in patients with addictive behaviors.^41^ Therefore, instead, the increased response time likely relates to delayed disengagement, prompted by a greater *PE* signal from the prior loss trial, rather than an allocation of cognitive resources to manage behavior in probabilistically riskier regimes. In the context of substance abuse disorders, these findings may relate to a predisposition of impulsive choosers to relapse, instead of opting for potential longer-term rewards.

### Reward Models and Encoding

While a signature of compulsive reward prediction has been discovered in the human nucleus accumbens,^61^ a notable focus of current research is *how* such reward prediction information is propagated into decision making circuits that mediate perception and action. Following the clear behavioral differences between impulsivity groups, we expected there to be differences for *RVE* and *RPE* variables in terms of TD model outcomes (i.e., learning rates) and neural encoding. Similar to previous reward TD learning work in fMRI,^40^ we observed value estimates approach an asymptote after approximately 40 trials, after which value estimations remain relatively stable (Figures 2a,e & 3a,e). We also see positive *RPEs* for rewarded trials and negative *RPE*s for unrewarded outcomes (Figures 2d,h & 3d,h).

For value expectation we saw that individuals with higher impulsive choice levels were seemingly influenced by their internal value representations, which they had built up over trials, rather than value expectation placed on the external cue (i.e., balloon color). This suggests that more impulsive choosers have a more rigid value expectation, which may support implementation of a more model-free learning paradigm.^62–64^ Recently, a reinforcement learning modeling approach has recognized a middle ground between model-free and model-based learning in addiction disorders.^65^ This successor representation modeling also highlighted rigidity of value expectations. Previous research has shown that individuals with addiction disorders tend to use successor representation of states, rather than model-free or model-based learning.^66^ Our results potentially align with such successor representation-like learning patterns, as MI choosers have stricter value representations than LI choosers, evident from maladaptive decision-making behaviors observed after an unrewarded trial.^66^ The value representation rigidity that we observe has also been highlighted in different task structures. Prior research using a reversal learning task showed that MI choosers exhibit increased perseveration (i.e., delayed disengagement) after the reversal learning point, around reward probabilities.^27^ Concurrent with our findings, these results highlight a behavioral dependency and impaired inhibitory control of updating value, which stifles disengagement from current reward value expectations, emphasizing the importance of value expectation as an indication of impulsive choice behavior.

We found marked differences in encoding of reward variables in the brain. Overall, both *RVE* and *RPE* were represented in MTG, MFG, HIPP, and ACgG. Between impulsivity groups, MI subjects had higher encoding in middle temporal, frontal, and hippocampal regions. Previous work also implicates hippocampal regions and surrounding mesial temporal regions to reward surprise and punishment avoidance.^67–69^ Therefore, it is unsurprising to see both value and prediction error encoded to a greater extent in these regions, especially greater *RPE* encoding. The dopaminergic error signal is key for economic decisions, by helping to modulate value expectation for upcoming trials,^70^ which may explain why there is more widespread neural representation of *RPE* than *RVE*, our error learning signal is updating the basal value expectation function. Additionally, in our asymmetric model encoding, we saw greater encoding of *positive RPE* by LI choosers. *Positive RPE,* but not *negative RPE*, was primarily encoded in the amygdala by both groups, a region that did not significantly encode signed *RPE*s in the original TD models.

Interestingly, MI choosers exhibited greater encoding of all reward variables, compared to LI choosers. *RVE* may be an important component of impulsive choice, where higher value encoding relates to clinically relevant increases in impulsivity.^71^ Furthermore, these findings support the idea of reward hypersensitivity in impulsive choosers, where a greater activation of these regions is indicative of a sensitivity toward reward and reward-oriented processes. Previous research has found greater amplitudes of the P2a event-related potential amplitude in OFC for self-reported high impulsive choosers, compared to low impulsive choosers, highlighting potential reward hypersensitivity.^72^ The current results expand on the focal reward sensitivity findings by examining reward encoding in localized correlates of population neuronal firing across the entire brain.^72,73^

Generally, reward was encoded in the left hemisphere, but we observed hemispheric differences in impulsivity groups, where MI choosers tended to encode reward model variables in the left hemisphere and LI individuals tended to encode reward model variables in the right hemisphere. Prior research found expected value to be represented across brain regions in a distributed fashion,^74^ but did not examine differences between MI and LI groups. Moreover, some prior studies have highlighted the left hemisphere as a common core of value representations in the brain, (specifically left VMPFC, left DLPFC, and left cerebellum)^75^ and as the hemisphere involved in encoding recall of high-value items encoding over low-value item.^76^ We speculate that the left hemisphere, having dominance, may be involved in impulsivity (i.e., cue-induced craving)^77^ and motor planning^78^ to a greater extent than the right hemisphere. It is possible that impulsive choosers, that have rigid value representations also encode value in a more lateralized fashion, isolated to the left hemisphere.

### Risk Models and Encoding

A separable but intertwined process in reinforcement learning with uncertain outcomes is risk estimation.^11,12,78^ Economic risk is generally defined as a quadratic function of reward uncertainty.^11^ Reasoning that risk estimation is an important element of decision making under outcome uncertainty, we sought to understand trial-by-trial updating of risk expectation and surprise using temporal difference learning models. Previous fMRI research has found discrete brain areas in the anterior insula and striatum that explicitly represent risk^11,12^ and risk prediction error.^79^ Furthermore, an fMRI study of BART found explicit risk signals in the dorsal ACC, anterior insula, and inferior frontal sulcus.^80^ These studies examine components of a similar risk processing network in the human brain to that which we uncovered using temporal difference learning models. Our risk model also highlighted risk processing in anterior insula and ACC by MI choosers, specifically for Risk expectation and both positive and negative *PE*s.

Several salient differences between reward and risk encoding in human neuronal population activity arose from our analyses. Risk variables were generally less widely represented than reward variables. In contrast to reward *VE* and *PE*, *Risk VE* was more widely represented than *Risk PE*. While *Risk VE* straightforwardly approximates the balloon color variable, the interpretation of *Risk PE* is less straightforward, yet crucial for learning reward probabilities and encoding risk in a stochastic environment. *Risk PE* represents surprise about the level of uncertainty around reward, which may reflect participants’ outcome-aligned actualized risk behavior. We saw that LI choosers tended to modulate *VE* after an error, but MI choosers were more rigid in their value estimates throughout the task, which demonstrates the inclination of MI choosers to over-anticipate reward but to not adapt behavior, accordingly. That *Risk PE* exhibited the smallest representation in the HFA signal may speak to the lack of utility in explicitly representing such a higher order variable in value-oriented neuronal computations under uncertainty. Neurally, LI subjects encoded *Risk PE* to a greater extent than MI subjects. It is possible that greater risk encoding allows for better representation of reward potential, as we see, behaviorally, greater overall points in the task compared to MI subjects. Across subjects, we found that MI choosers were more likely to have significantly different response times in response to the risk cue, but, interestingly, more likely to have a negative correlation between previous trial risk prediction and response time on the current trial. We interpret this across-subject result to reflect a lack of delayed discounting in response to risk cues in MI choosers and an inversion of risk estimation between LI and MI choosers.^81–83^

Similar to reward, we see hemispheric differences between impulsivity groups, where MI choosers have greater encoding of risk in the left hemisphere, whereas LI choosers have greater encoding of risk in the right hemisphere. As our subject cohort has drug-resistant epilepsy, the electrode contacts are placed based solely on clinical considerations. It is common for lateralized epileptiform activity to originate in the left temporal lobe.^84^ Although the majority of clinical sEEG electrodes are placed in the left hemisphere this does not explain our hemispheric differences findings as our proportion analysis accounts for uneven contact numbers in each hemisphere.

## Limitations and Future Directions

We endeavored to achieve rigorous data collection and meticulous analysis of the data for this study. However, there are some inherent limitations which we address here. First, the electrodes that we are able to record from are placed with clinical consideration, which limits potential recording sites of interest, from a research perspective. It is possible that there are regions encoding reward and risk variables that we are not able to sample. That being said, we have numerous electrodes in cortical, mesial temporal, and deep brain structures from which to infer reward and risk encoding. Second, while we try to make the study engaging, it is possible, as with any computer-based cognitive paradigm, that subjects could be responding quickly, and therefore seemingly more impulsively, due to amotivation. To minimize the potential for amotivation, we introduce the task goals to the subjects prior to task initiation (see Methods for task script). Furthermore, we ran across-subjects analysis on the first 50 trials and last 50 trials to examine learned changes in IT throughout the task. If subjects were becoming less motivated, we would expect a decrease in inflation time toward the end of the task and that this effect would be exacerbated by the more impulsive choosers who are banking point earlier. However, we see a significant increase in inflation time at the end of the task compared to the start of the task (first 50 trials inflation time; *M* = 3.92s ±1.80; last 50 trials inflation time; *M* = 4.12s ±1.99; *t*(44) =-2.18, *p* = 0.035). We can therefore be confident that the subjects are engaged and learning throughout the task (see Supplementary Figure 10).

While we show activation of different brain regions related to reward and risk, we do not show the how the flow of information through the neural circuits is occurring. One future direction is to complete this analysis, which means that we can show the implications of impulsivity and reward in the brain.

## Conclusion

Here, we highlight anatomical differences regarding the neural processes of reward and risk, modulated by impulsivity. This large dataset of direct electrical recordings from the human brain offered a unique opportunity to analyze the neural underpinnings of impulsivity across subjects. We detail the patterns of neuronal population activity and brain areas involved with impulsive choosing. Understanding these neural associations sheds light on the brain mechanisms underlying impulsivity disorders and relapse in substance use, and therefore may lead to improved or more relevant biomarkers for neural prostheses. Utilization of these biomarkers to detect impending impulsive choices lead to novel, targeted strategies for treating addiction disorders.

## Methods

### Experimental Model and Subject Details

#### Subjects

Research participants were drug-resistant epilepsy patients (21 female, *M* = 36 ± 10 years of age) who were undergoing intracranial monitoring as part of neurosurgical treatment for their seizures. Please refer to the results section of the main manuscript for additional information related to the study population (e.g., sample size). Full details on gender and age for all study participants can be found in the supplementary materials (Table S3). All participants provided informed consent prior to any data collection. The University of Utah Institutional Review Board approved this study.

### Method Details

#### Inclusion/Exclusion

Potential research participants were drawn from a population of patients slated to undergo intracranial neuromonitoring for drug resistant epilepsy. Participants were recruited prior to their stay in the epilepsy monitoring unit (EMU) at the University of Utah. Two participants were excluded from analysis, having been unable to complete enough task trials to achieve sufficient across-trial statistical power.

#### Balloon Analogue Risk Task (BART)

BART is a cognitive paradigm that can measure both impulsivity and risk-taking behaviors by conceptualizing the probability for potential for reward.^85^ The aim of BART is to inflate a simulated balloon without the balloon popping in order to gain points that are linearly related to the size of the balloon and the inflation time (IT). If the balloon pops the participant neither wins or loses points. For each trial, balloon inflation is initiated by a button press and stopped by same button press. There are two types of trials: 1) passive, in which a colored balloon will inflate to the size of a gray circle threshold and the subjects will always gain points and 2) active, where the subjects must actively stop the balloon before it pops, but try to gain as many points as possible, which corresponds to the IT duration. For both active and passive trials, there are three colors categories of balloon: yellow, orange, and red, which have greater risk levels related to IT, respectively. To stop subjects learning exact inflation sizes, each balloon has a randomized inflation range calculated from a normal distribution and standard deviation, in which the balloon can pop at any timepoint. A balloon (yellow, orange, or red) appears on the screen, so the subjects can assess the relative risk of the balloon before pressing a button to start the balloon inflation, then pressing a button to stop the inflation, before the balloon pops. Overall, the larger the balloon IT, the greater risk level the participants are willing to take, while the earlier the participants bank the points, indicates a greater level of impulsivity (i.e., favoring smaller certain reward over larger less certain reward).

Before starting BART, participants were given instructions on how to complete the task. Here is an example text:

> *“The overall aim of the task is to inflate the balloons to gain as many points as you can. The bigger the balloon gets the more points you will get, but, if you leave the balloon to inflate for too long, it will pop and you won’t get any points. There are also different colors of balloons: red balloons inflate the least before popping, orange inflate to an intermediate level before popping, and yellow balloons inflate the most before popping. There are also grey trials where you get no points and passive trials where you can learn about how big the different colors of balloons can get and get points for free. Both of these types of balloons will appear with a gray circle indicating the size to which the balloon will inflate.”*

#### Data Collection

Each participant carried out the BART while undergoing monitoring in the EMU. The task was implemented in PsychToolbox for Matlab.^86,87^ Participants registered their behavioral responses via a USB game controller (Logitech G F310). A typical task session began with a researcher administering the aforementioned verbal instructions to the participant. The participant then completed over 200 trials of BART while we recorded electrical activity from intracranial stereoelectroencephalographic depth (sEEG) or subdural electrocorticographic (ECoG) electrodes implanted solely for localization of epileptogenic tissue.

Two participants included in this dataset did BART twice. As there were no pre-allocated across-subject study conditions, the task was neither randomized nor blinded at the level of research participants. The order in which task trials were presented was randomly generated.

#### Electrophysiological Recordings

Signals from clinical electrodes were recorded during task sessions at 1000 samples per second at 16-bit resolution on a 128-channel neural signal processor (Blackrock Microsystems, Salt Lake City, UT). The signal was reconstructed without its first principal component across time in order to remove any reference or line noise artifacts. Task events and patient behavior were synchronized with the neural recording at sub-millisecond precision via digital codes sent from the psychtoolbox code to a PCI-express card in the task control computer. Trials and channels with interictal epileptiform discharges were excluded by detecting outliers in the range of voltage signals recorded on each channel.

#### Anatomical Localization

Electrodes were localized using the LeGUI software package.^88^ Briefly, preoperative MRIs were co-registered to post-implant CTs, and electrodes were detected automatically via a density threshold. All automatic localizations were verified manually. LeGUI fits these localized electrodes into a standard space (Montreal Neurological institute 152; MNI), from which the Neuro Morphometric atlas anatomical locations were derived (Neuromorphometrics, Inc.) Proportion tests were carried out on these categorical anatomical labels with a significance criterion of 0.05.

Flat cortical surface representations were generated using PyCortex.^89^ To do this, we first generated three-dimensional models of each hemisphere of the MNI brain using freesurfer.^90^ We then removed the medial surface around the corpus callosum and made five cuts along the mesial aspect of the inflated pial surface. Only electrodes within 6 mm of the pial surface were retained. Those electrode locations were projected onto the nearest vertex of the three-dimensional surface. The cortical surface from each hemisphere was then flattened using PyCortex. An anatomical guide to the flat brain representations is shown in Figure S4.

### Quantification and Statistical Analysis

#### Impulsivity Metric

Optimal BART performance involves inflating balloons as much as possible, while minimizing balloon popping. Impulsive choice in BART is characterized by stopping balloon inflation early in order to avoid risk and gain a smaller, more immediate reward. We therefore operationally defined impulsive choice as the distributional similarity between active and passive IT distributions. Inspired by active inference, a log Kullback-Leibler Divergence (KLD) was used to measure distributional similarity between active and passive trial IT distributions.^91^ We also corroborated this measure via proximity of active and passive trial IT distribution means using a *Z*-statistic (Figure S3b). For BART, IT distribution for passive trials was used as the reference distribution. For example, more similar active and passive trial distributions reflected a lower impulsive choice score as the participant allowed balloons to inflate to similar sizes as the passive trials, maximizing reward at the cost of accuracy. Alternatively, more impulsive choices reflected consistently shorter active IT distributions and higher impulsive choice scores. A one-dimensional gaussian mixture model of the KLD-derived impulsivity scores was used to classify subjects into more impulsive or less impulsive choosers (see Manuscript Figure 1f). ANOVA was used to test for differences in accuracy across balloon colors between categories of impulsive choosers. A Mann-Whitney U test was used to test for score differences between impulsivity categories. Ordinary linear regression with ANOVA was used to test for systematic relationships between KLD-derived impulsivity levels and both accuracy and overall reward (Figure 1I-J). Significance criteria was set at 0.05 for these tests.

#### Temporal Difference (TD) Learning Models

In order to infer the neural underpinnings of reward and risk expectation and surprise, we fit discrete-time reinforcement learning models to patients’ behavior using maximum likelihood estimation.^43,92^ For each trial, *t*, a value estimate (*VE*) was determined based on the *VE* from the previous trial and the current trial’s prediction error (*PE*), scaled by the learning rate (α):

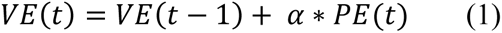

Where *PE* on each trial was defined as the difference between the previous trial’s *VE* and the reward received, *R*:

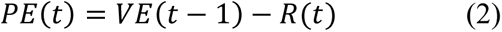

VE for reward and risk models were initialized to 0 and 0.5, respectively. The value of R for risk models was the cumulative reward probability on each trial. We fit asymptotic functions to VE estimates across trials using the Matlab function ‘fit’ with coefficients 1 and 1/x (red lines in Figures 2e & 3e).

Optimal learning rates were derived from maximum inverse temperatures, which were estimated from outcomes using the *softmax* rule for outcome probability on each trial:^43^

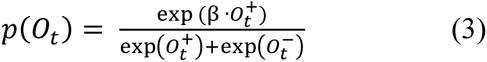

Where β is the inverse temperature, and *O*_*t*_ refers to outcome categories on each trial. We tested for any association between optimal learning rates and impulsivity scores using Mann Whitney U tests with a significance criterion of 0.05.

#### Neural correlates of TD model variables

To understand which brain areas encoded these temporal learning model variables across trials, we examined broadband high frequency (HFA: 70 – 150Hz) local field potentials recorded from intracranial electrodes. HFA is an established correlate population neuronal firing near the electrode.^46,93,94^ HFA was isolated by filtering the voltage signal from each channel between 70 and 150 Hz using a 4^th^ order Butterworth filter. The Hilbert transform of the filtered signal was smoothed with a 50 ms moving window and aligned on the appearance of the balloon (for risk models), and on outcome (for reward models). We modeled each subject’s HFA on each trial as a linear function of the TD model variables (value and prediction error) for both risk and reward elements of BART:

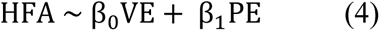

Where the β values represent the regression coefficients for HFA responses to risk and reward expectation (i.e., *VE*) and surprise (i.e., *PE*). The risk model data came from cue-aligned responses (i.e., responses to the appearance and color of the balloon, which signaled each balloon’s relative risk; Figure 1), and the reward model data came from outcome-aligned responses (i.e., points gained, or balloon popped). One second of HFA data, 250 ms after the aforementioned events was included in the models. The *VE* and *PE* variables were taken from the TD model with the optimal learning rate.

The balloon pop is a relatively salient sensory element of the task. In order to control for the salience of this stimulus, we fit additional generalized linear mixed effects models that controlled for brain areas that may be responsive to the sensory salience of the balloon pop:

##### (A) Salience controlled TD Models

HFA ∼ ValueEstimate + RewardPE + (ValueEstimate|outcome) + (RewardPE|outcome)

HFA ∼ RiskEstimate + RiskPE + (RiskEstimate|outcome) + (RiskPE|outcome)

Moreover, we were interested in understanding neuronal population encoding of different types of prediction errors. Prediction error is signed signal, unsigned prediction error is the absolute value of the magnitude of the prediction error signal. Positive prediction error is the rectified signal and negative prediction error if the rectified signal in the negative direction. We therefore fit brain data to both unsigned and asymmetric prediction errors via the following models:

##### (B) Salience controlled Unsigned TD Models

HFA ∼ ValueEstimate+UnsignedRewardPE + (ValueEstimate|outcome) +(UnsignedRewardPE|outcome)

HFA ∼ RiskEstimate + UnsignedRiskPE +(RiskEstimate|outcome) + (UnsignedRiskPE|outcome)

##### (C) Salience controlled Asymmetric TD Models

HFA ∼ ValueEstimate + positiveRewardPE + negativeRewardPE + (ValueEstimate|outcome) + (positiveRewardPE|outcome) + (negativeRewardPE|outcome)

HFA ∼ RiskEstimate + positiveRiskPE + negativeRiskPE + (RiskEstimate|outcome) + (positiveRiskPE|outcome) + (negativeRiskPE|outcome)

#### Proportion Tests

We sought to understand which brain regions across patients and contacts significantly encoded TD model variables. To test for significant proportions of TD variable encoding contacts in particular brain regions, we carried out a series of proportion tests on the NMM anatomical labels across recording contacts, with significance criteria set to 0.05. First we tested for significant proportions of contacts over the three impulsivity categories (less, more, both) and then over the significant model coefficients (reward expectation, reward surprise, risk expectation, risk surprise, reward and risk interactions, and all contacts). We also tested for hemispheric differences in TD variable encoding using these NMM anatomical labels.

#### Post-Outcome Response Time Analysis

We looked to examine how response time tendencies aligned with reward and risk TD model variables. Therefore, we examined RT differences between MI and LI choosers relative to two TD model variables: cue-aligned value expectation on the current trial and prediction error from the previous trial. To do this, we first took a section of *VE*s and *PE*s that were similar based on their median and one standard deviation. For the VE analysis, we wanted to see if the cue-aligned *VE*s predicted RTs between impulsivity groups. We regressed the correlation coefficient of an ANOVA model *VE* and RTs amongst balloon colors against impulsivity level. As an additional control, we regressed the *Risk PE* and RT *t*-value on the previous trial against impulsivity level (see Supplementary Figures S8 & S9).

### Key Resources Table

| REAGENT or RESOURCE | SOURCE | IDENTIFIER |
| --- | --- | --- |
| Software and Algorithms |  |  |
| MATLAB | Mathworks | https://www.mathworks.com |
| Blackrock Recording | Blackrock Neurotech. | https://blackrockneurotech.com |
| Psychtoolbox | Psychtoolbox | http://psychtoolbox.org |
| LeGUI | Mathworks | https://github.com/Rolston-Lab/LeGUI |
| Pycortex | Python | https://www.python.org/ |
| FreeSurfer | Python | https://www.python.org/ |
| PCI-e cards | National Instruments | https://www.ni.com/ |
| BNC breakout boards | National Instruments | https://www.ni.com/ |
| NMM Atlas | Neuromorphometrics, Inc. | https://www.neuromorphometrics.com/ |

**Table 1.** Percentage and raw proportions of significantly encoded contacts for LI, MI, and both impulsivity groups. NA highlights non-significant regions. All Reward and Risk model variables are included in the table.

| TD Variable | Encoding Regions<br>Less Impulsive Group |  |  |  |  |  |  | Encoding Regions<br>More Impulsive Group |  |  |  |  |  |  | Encoding Regions<br>Both Impulsive Groups |  |  |  |  |  |
| --- | --- | --- | --- | --- | --- | --- | --- | --- | --- | --- | --- | --- | --- | --- | --- | --- | --- | --- | --- | --- |
|  | MTG | MFG | HIPP | FuG | ACgG | ENT | MOrG | MTG | MFG | HIPP | ACgG | Ains | MOrG | SFG | MTG | MFG | HIPP | ACgG | FuG | MORG |
| RVE | 20<br>(6%) | 17<br>(9%) | 16<br>(9%) | 8 (9%) | 8 (9%) | NA | NA | 9 (4%) | 10<br>(7%) | 9 (6%) | 3 (4%) | 4<br>(4%) | 3<br>(7%) | NA | 45<br>(9%) | 43<br>(13%) | 45<br>(14%) | 17<br>(11%) | NA | NA |
| RPE | 106<br>(33%) | 55<br>(28%) | 52<br>(30%) | 29<br>(33%) | 27<br>(32%) | NA | NA | 63<br>(31%) | 43<br>(30%) | 52<br>(37%) | NA | NA | NA | NA | 169<br>(32%) | 98<br>(29%) | 104<br>(33%) | NA | NA | NA |
| RVE & RPE | 114<br>(36%) | 59<br>(31%) | 58<br>(33%) | 30<br>(34%) | 32<br>(38%) | 32<br>(44%) | NA | 69<br>(33%) | 44<br>(31%) | 58<br>(41%) | NA | NA | NA | NA | 183<br>(35%) | 103<br>(31%) | 116<br>(37%) | 50<br>(32%) | NA | NA |
| Risk VE | 12<br>(4%) | 26<br>(13%) | 17<br>(10%) | 10<br>(11%) | 11<br>(13%) | 9<br>(12%) | 26<br>(43%) | 9 (4%) | 10<br>(7%) | 9 (6%) | 3 (4%) | 4<br>(4%) | 3<br>(7%) | NA | 21<br>(4%) | 36<br>(11%) | 26<br>(8%) | 14<br>(14%) | 12<br>(11%) | 13<br>(12%) |
| Risk PE | 22<br>(7%) | 28<br>(15%) | 15<br>(9%) | NA | 8 (9%) | NA | 9<br>(15%) | 9 (4%) | 15<br>(11%) | 9 (6%) | 14<br>(20%) | NA | NA | 4<br>(12%) | 31<br>(6%) | 43<br>(13%) | 24<br>(8%) | 22<br>(14%) | NA | NA |
| Risk VE &<br>Risk PE | 33<br>(10%) | 43<br>(22%) | 31<br>(18%) | 16<br>(18%) | 15<br>(18%) | NA | 19<br>(32%) | 17<br>(8%) | 20<br>(14%) | 14<br>(10%) | 15<br>(21%) | 7<br>(7%) | NA | NA | 50<br>(10%) | 63<br>(19%) | 45<br>(14%) | 30<br>(19%) | 19<br>(17%) | 24<br>(23%) |
| Total<br>Contacts | 318 | 193 | 176 | 88 | 85 | 73 | 60 | 206 | 142 | 140 | 70 | 51 | 43 | 33 | 524 | 335 | 316 | 155 | 114 | 105 |

## Acknowledgements

We would like to acknowledge the Neurosurgery Department at the University of Utah. Thank you to our funding sources at the NIH for making this work possible. NIH: R01MH128187

## Author Contributions

Conceptualization, E.H.S. and R.L.C.; Methodology, E.H.S., R.L.C., T.D., B.K., S.R., and J.D.R.; Investigation, E.H.S., and R.L.C., T.D., B.K., S.R., and J.D.R.; Writing – Original Draft, R.L.C.; Writing – Review & Editing, R.L.C., E.H.S., T.D., B.K., S.R., and J.D.R.; Funding Acquisition, E.H.S.; Resources, E.H.S. and J.D.R.; Supervision, E.H.S.

