## Supplementary Materials for "More widespread and rigid neuronal representation of reward expectation underlies impulsive choices"

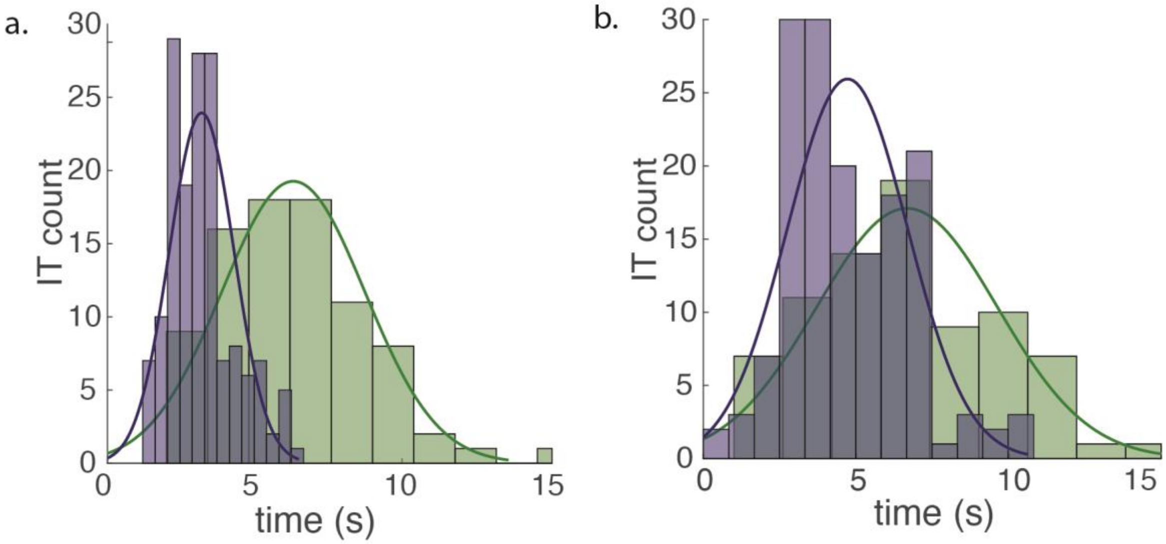


**Figure S1. Balloon Inflation Time Distributions. (a)** More Impulsive Individual example (KLD score: 2.54) balloon distribution inflation times for passive (green) trials (inflation time *M*: 6.31s) and active (purple) trials (inflation time *M*: 3.2s). **(b)** Less Impulsive Individual example (KLD score: 0.43) balloon distribution inflation times for passive (green) trials (inflation time *M*: 6.66s) and active (purple) trials (inflation time *M*: 4.71s).

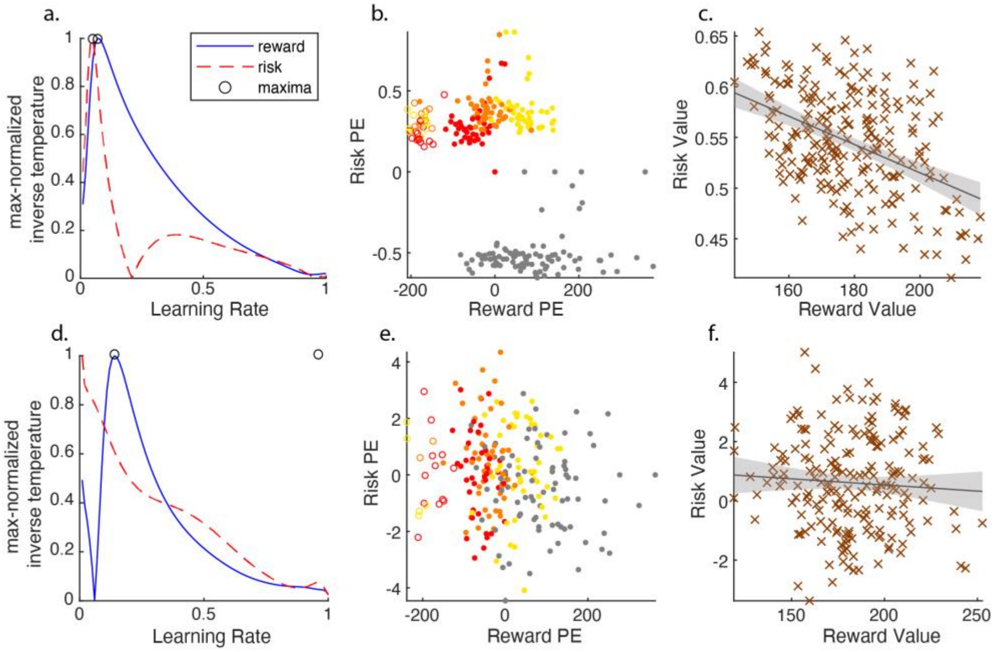


**Figure S2. Examples of Temporal Difference Models and Optimal Alphas in MI and LI Individuals. a-c** (KLD: 1.0718), $t(215) = -7.36$, $p = 0.00$, $R^2 = 0.20$. **d-f.** (KLD: 2.6961) $t(195) = -0.91$, $p = 0.37$, $R^2 = 0.00$.

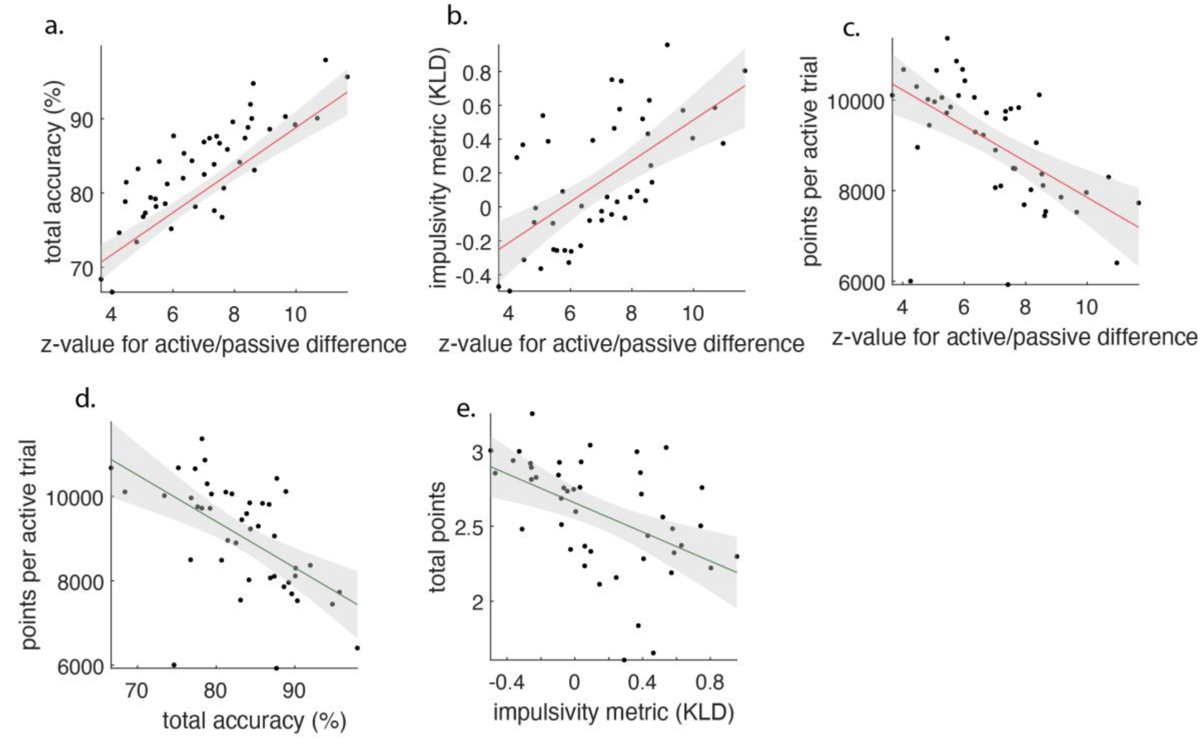


**Figure S3. Impulsivity and Task Performance Regression Plots. (a)** Total task accuracy regressed against the z-value for active and passive trial IT differences. **(b)** Impulsivity scores regressed against the z-value for active and passive trial IT differences (adjusted-$R^2$ = 0.38, $F(2, 45)$ = 27.50, $p = 4.51 \times 10^{-6}$). **(c)** Total points for active trials regressed against the z-value for active and passive trial *IT* differences. **(d)** Total points for active trials regressed against total task accuracy. **(e)** Total points regressed against impulsivity scores.

**Table S1. Formulas for Temporal Difference, Unsigned, and Asymmetric Models.** TD model variables reward and risk.

<table>
<tr><td>TD: Reward</td><td>HFA ~ ValueEstimate + RewardPE + (ValueEstimate|Outcome) + (RewardPE|Outcome)</td></tr>
<tr><td>TD: Risk</td><td>HFA ~ RiskEstimate + RiskPE + (RiskEstimate|Outcome) + (RiskPE|Cue)</td></tr>
<tr><td>TD Unsigned: Reward</td><td>HFA ~ ValueEstimate + UnsignedRewardPE + (ValueEstimate|Outcome) + (UnsignedRewardPE|Outcome)</td></tr>
<tr><td>TD Unsigned: Risk</td><td>HFA ~ RiskEstimate + UnsignedRiskPE + (RiskEstimate|Outcome) + (UnsignedRiskPE|Cue)</td></tr>
<tr><td>TD Asymmetric: Reward</td><td>HFA ~ RewardEstimate + positiveRewardPE + negativeRewardPE + (RewardEstimate|Outcome) + (positiveRewardPE|Outcome) + (negativeRewardPE|Outcome)</td></tr>
<tr><td>TD Asymmetric: Risk</td><td>HFA ~ RiskEstimate + positiveRiskPE + negativeRiskPE + (RiskEstimate|Outcome) + (positiveRiskPE|Cue) + (negativeRiskPE|Cue)</td></tr>
</table>

**Table S2. Task Performance.** Inflation Times and Response Times Between Impulsivity Groups.

| Group | Free ITs (s) ± SD | Control ITs (s) ± SD | Banked ITs (s) ± SD | Popped ITs (s) ± SD | Red ITs (s) ± SD *All* | Red ITs (s) ± SD *Banked* | Red ITs (s) ± SD *Popped* | Orange ITs (s) ± SD *All* | Orange ITs (s) ± SD *Banked* | Orange ITs (s) ± SD *Popped* | Yellow Balloon ITs (s) ± SD *All* | Yellow Balloon ITs (s) ± SD *Banked* | Yellow Balloon ITs (s) ± SD *Popped* |
|---|---|---|---|---|---|---|---|---|---|---|---|---|---|
| More Impulsive | 3.69 ± 0.71 | 6.77 ± 0.25 | 3.64 ± 1.84 | 3.64 ± 1.99 | 2.52 ± 0.88 | 2.38 ± 0.81 | 2.72 ± 0.93 | 3.25 ± 1.33 | 3.14 ± 1.26 | 4.19 ± 1.55 | 5.13 ± 2.08 | 4.98 ± 1.97 | 6.12 ± 2.44 |
| Less Impulsive | 4.33 ± 0.49 | 6.68 ± 0.2 | 4.42 ± 2.17 | 4.07 ± 2.22 | 2.68 ± 0.89 | 2.61 ± 0.85 | 2.75 ± 0.93 | 3.81 ± 1.33 | 3.69 ± 1.28 | 4.39 ± 1.40 | 6.44 ± 2.06 | 6.36 ± 2.0 | 6.77 ± 2.26 |

| Group | Free RTs (s) ± SD | Control RTs (s) ± SD | Banked RTs (s) ± SD | Popped RTs (s) ± SD | Red RTs (s) ± SD | Orange RTs (s) ± SD | Yellow RTs (s) ± SD | Post-Bank RTs (s) ± SD | Post-Pop RTs (s) ± SD |
|---|---|---|---|---|---|---|---|---|---|
| More Impulsive | 0.95 ± 0.34 | 0.91 ± 0.29 | 0.95 ± 0.34 | 97 ± 0.36 | 0.95 ± 0.36 | 0.94 ± 0.33 | 0.95 ± 0.33 | 0.93 ± 0.30 | 0.99 ± 0.45 |
| Less Impulsive | 0.83 ± 0.17 | 0.83 ± 0.17 | 0.83 ± 0.17 | 0.86 ± 0.20 | 0.82 ± 0.16 | 0.83 ± 0.18 | 0.84 ± 0.18 | 0.85 ± 0.17 | 0.78 ± 0.18 |

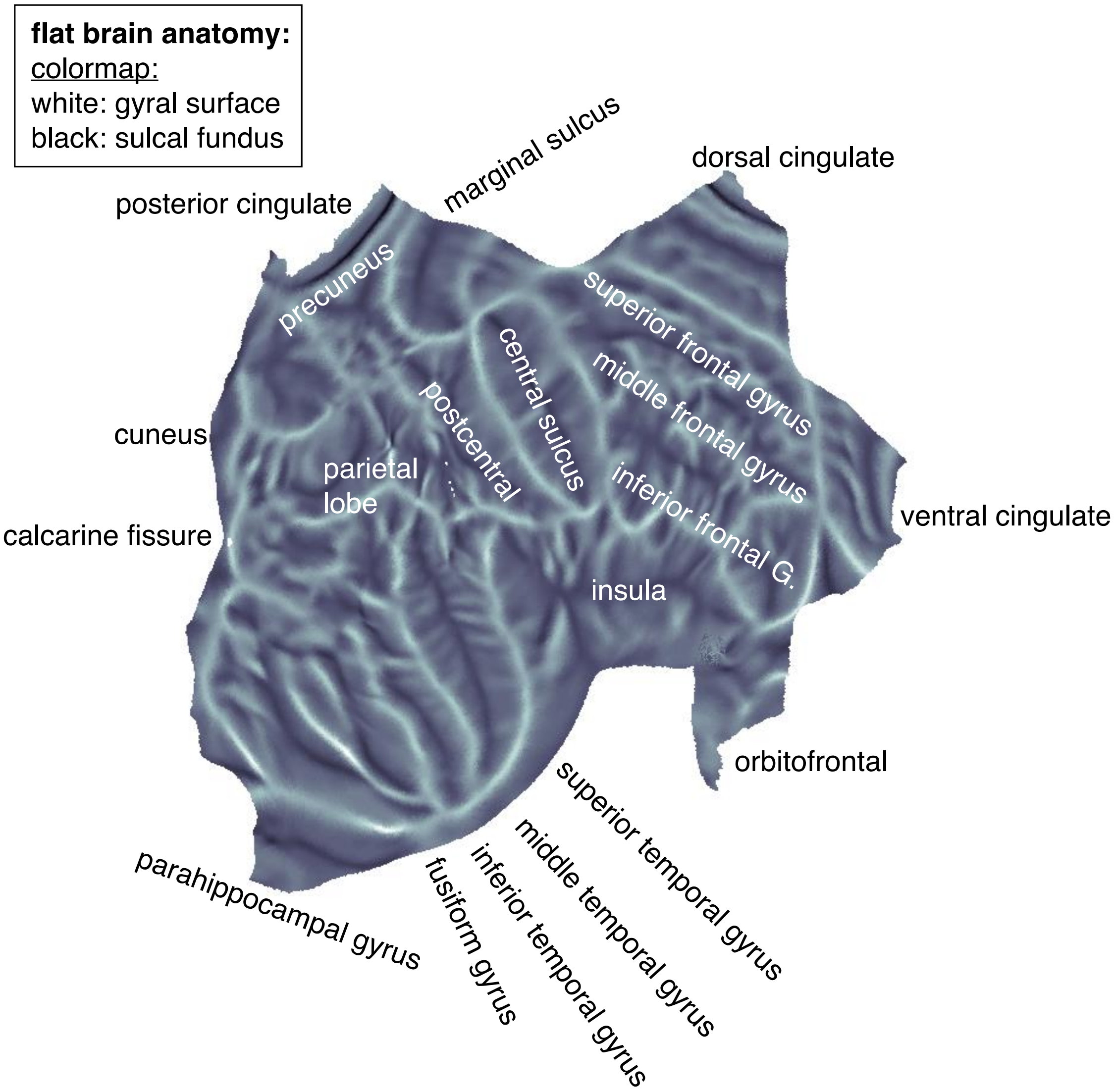


**Figure S4. Flat brain anatomy.** For flat brain plot references.

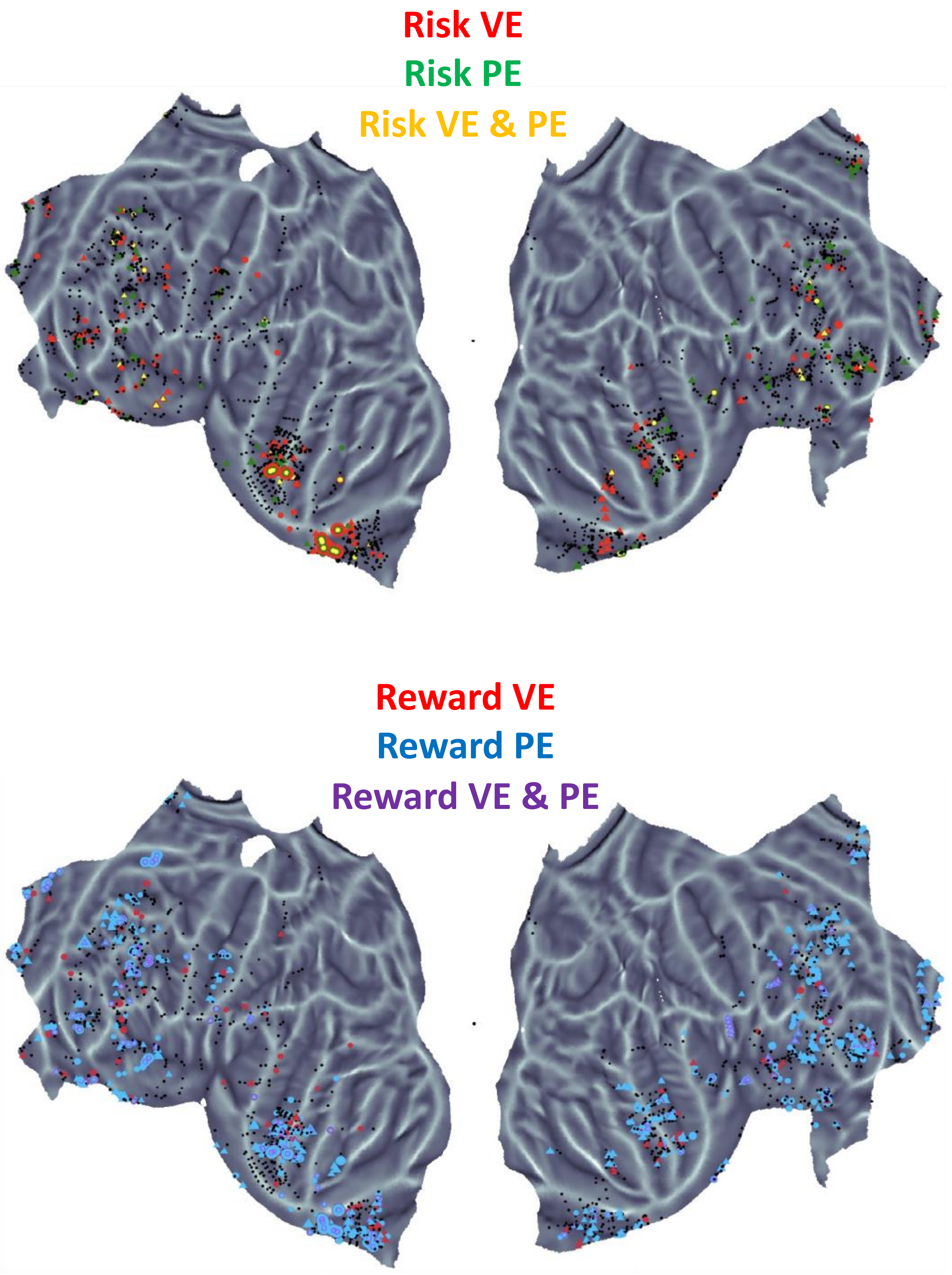


**Figure S5a. All Contacts Flat Brain Projections.** Risk VE (red), Risk PE (green), and Risk VE & PE (yellow) significantly encoded electrode contacts on flat brain. RVE (red), RPE (blue), and RVE & RPE (purple) significantly encoded electrode contacts on flat brain. This model includes all contacts.

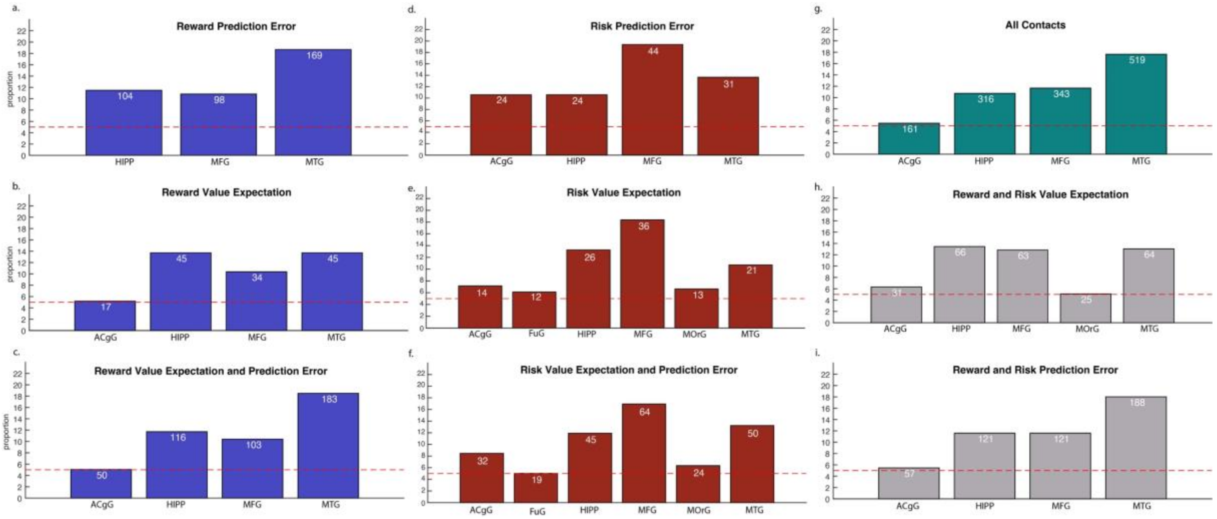


**Figure S5b. All Contacts Flat Brain: Proportion tests for Both Impulsivity Groups. (a)** Reward Prediction Error. **(b)** Reward Value Expectation. **(c)** Reward Value Expectation and Prediction Error. **(d)** Risk Prediction Error. **(e)** Risk Value Expectation. **(f)** Risk Value Expectation and Prediction Error. **(g)** All Risk and Reward Contacts. **(h)** Reward and Risk Value Expectation. **(i)** Reward and Risk Prediction Error.

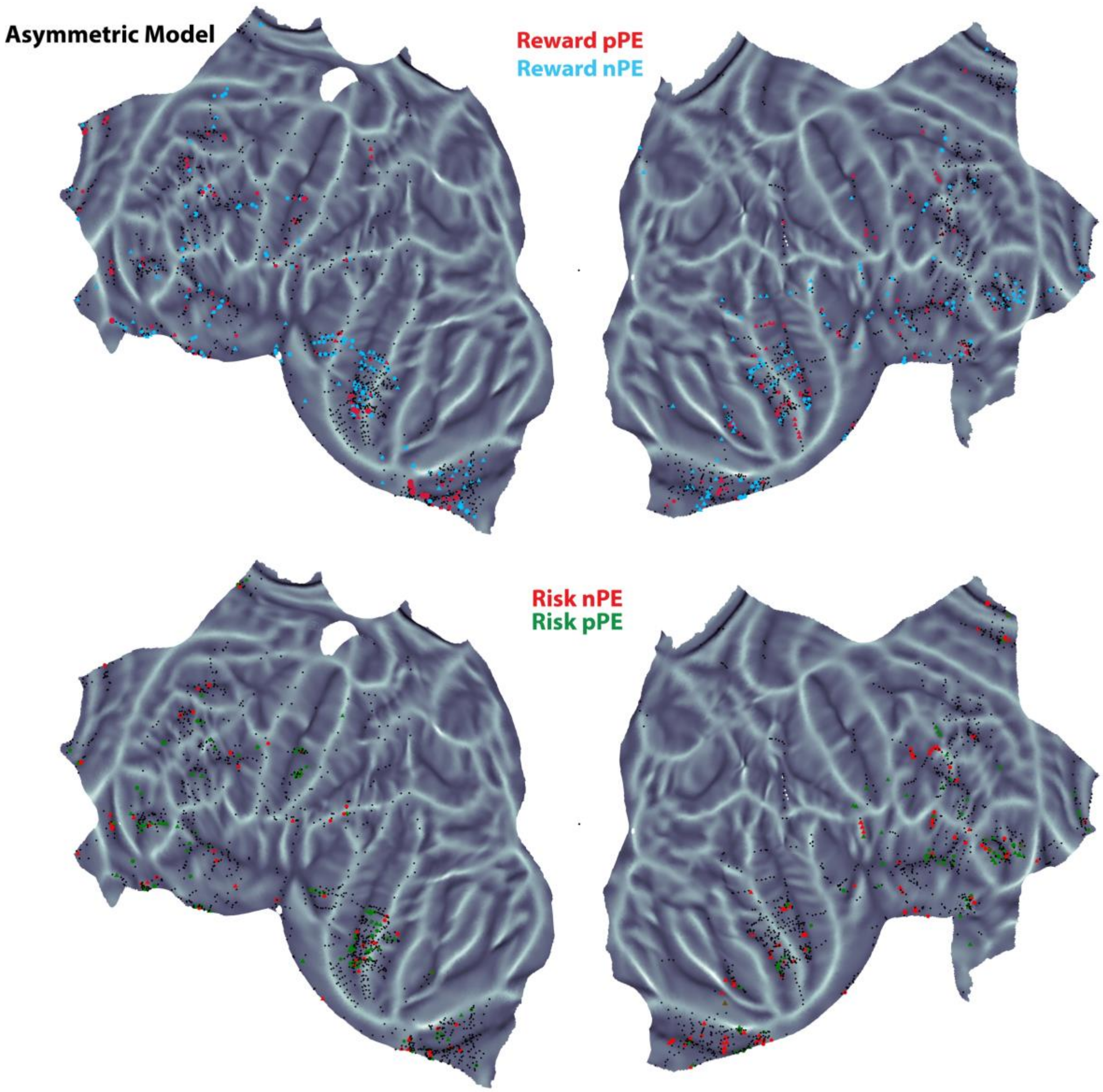


**Figure S6a. Asymmetric Model Flat Brain Projections. (a)** Reward positive PE (red) and Reward negative PE (blue) significantly encoded electrode contacts on flat brain, **(b)** Risk positive PE (red) and Risk negative PE (green) significantly encoded electrode contacts on flat brain.

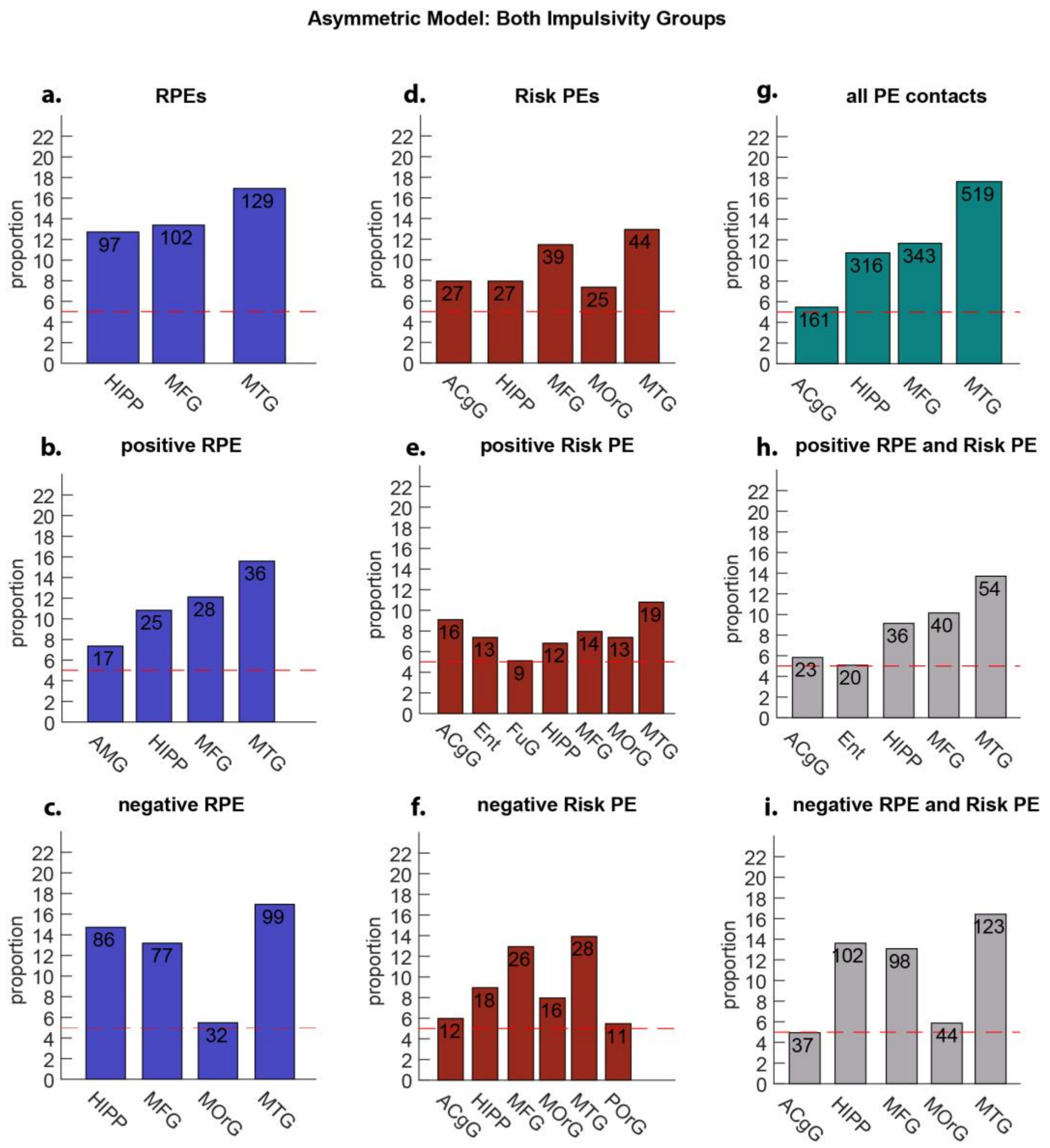


**Figure S6b. Asymmetric Flat Brain: Proportion tests for Both Impulsivity Groups. (a)** Reward Prediction Error. **(b)** Reward Positive Prediction Error. **(c)** Reward Negative Prediction Error. **(d)** Risk Prediction Error. **(e)** Risk Positive Prediction Error. **(f)** Risk Negative Prediction Error. **(g)** All Prediction Error Contacts. **(h)** Reward and Risk Positive Prediction Errors. **(i)** Reward and Risk Negative Prediction Errors.

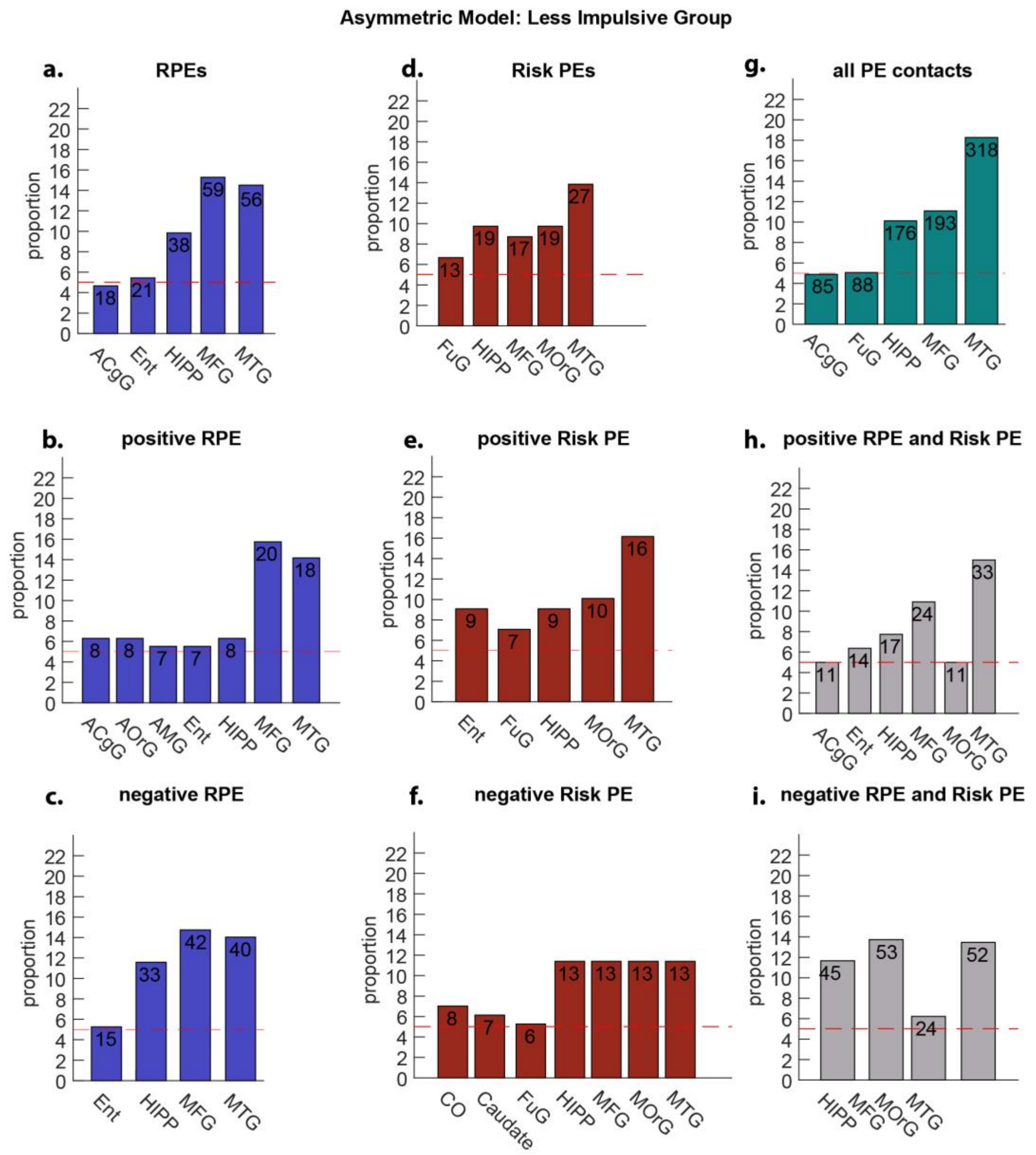


**Figure S6c. Asymmetric Flat Brain: Proportion tests for Less Impulsivity Groups. (a)** Reward Prediction Error. **(b)** Reward Positive Prediction Error. **(c)** Reward Negative Prediction Error. **(d)** Risk Prediction Error. **(e)** Risk Positive Prediction Error. **(f)** Risk Negative Prediction Error. **(g)** All Prediction Error Contacts. **(h)** Reward and Risk Positive Prediction Errors. **(i)** Reward and Risk Negative Prediction Errors.

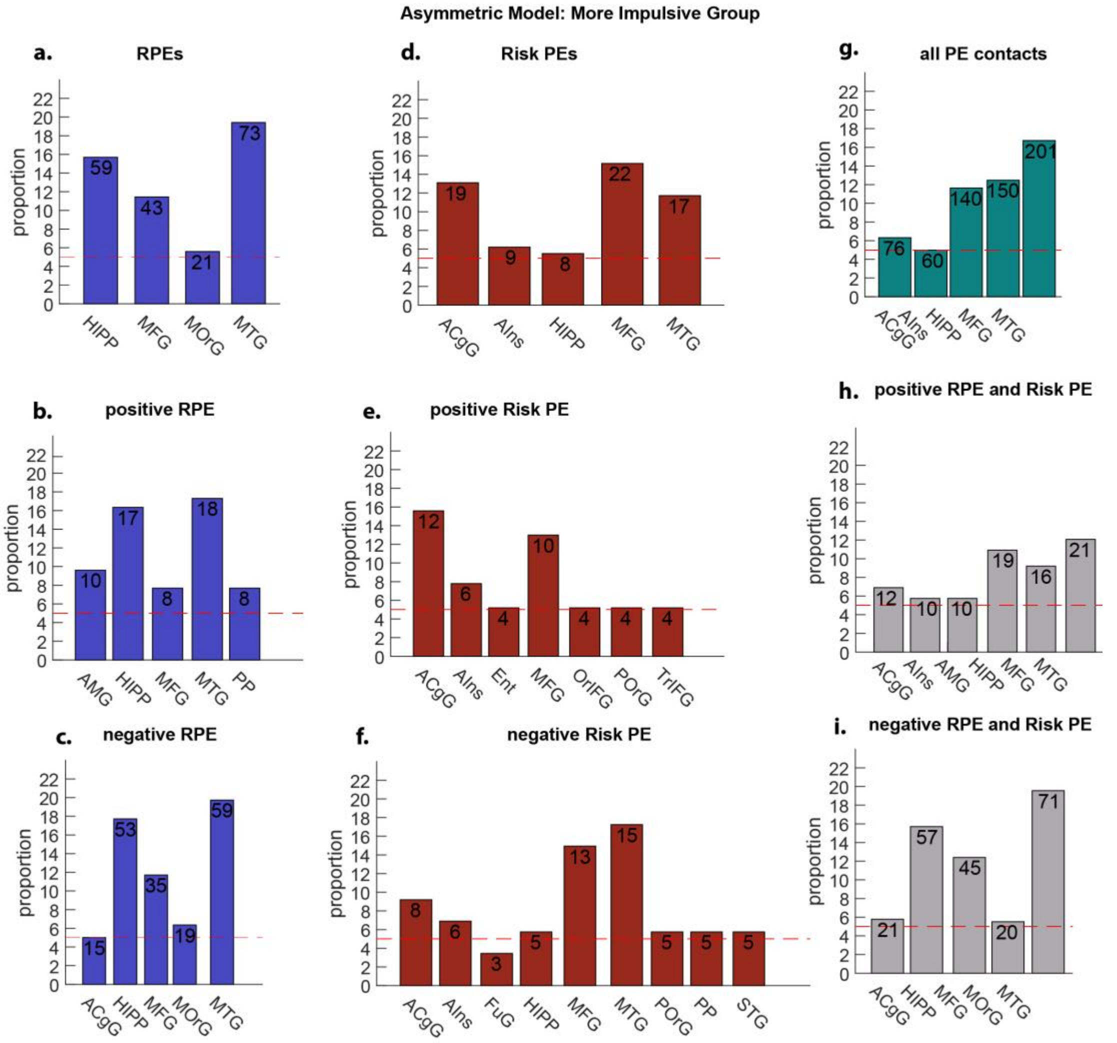


**Figure S6d. Asymmetric Flat Brain: Proportion tests for Less Impulsivity Groups. (a)** Reward Prediction Error. **(b)** Reward Positive Prediction Error. **(c)** Reward Negative Prediction Error. **(d)** Risk Prediction Error. **(e)** Risk Positive Prediction Error. **(f)** Risk Negative Prediction Error. **(g)** All Prediction Error Contacts. **(h)** Reward and Risk Positive Prediction Errors. **(i)** Reward and Risk Negative Prediction Errors.

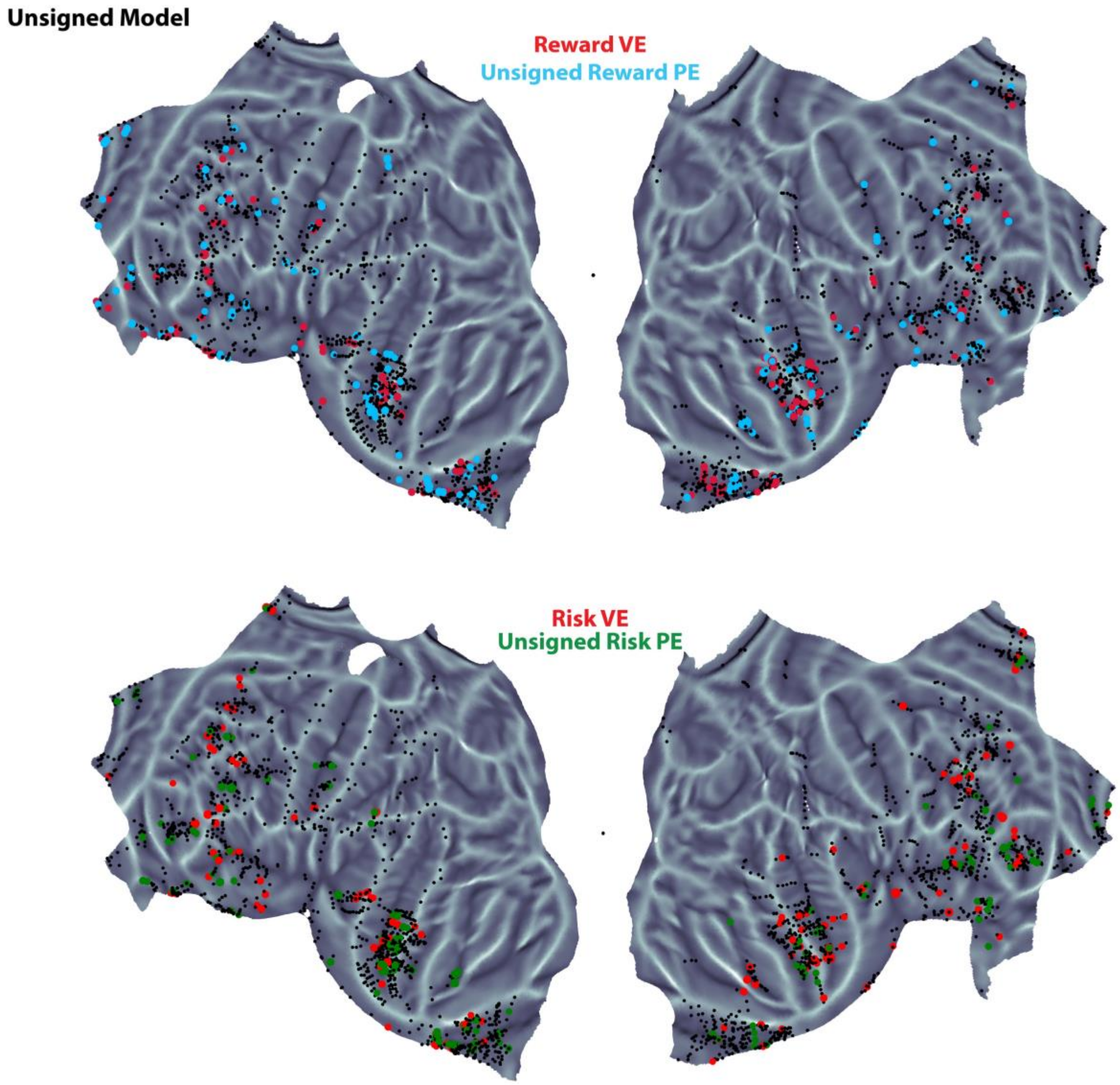


**Figure S7a. Unsigned Model Flat Brain Projections. (a)** RVE (red) and unsigned RPE (blue) significantly encoded electrode contacts on flat brain, **(b)** Risk VE (red) and unsigned Risk PE (green) significantly encoded electrode contacts on flat brain.

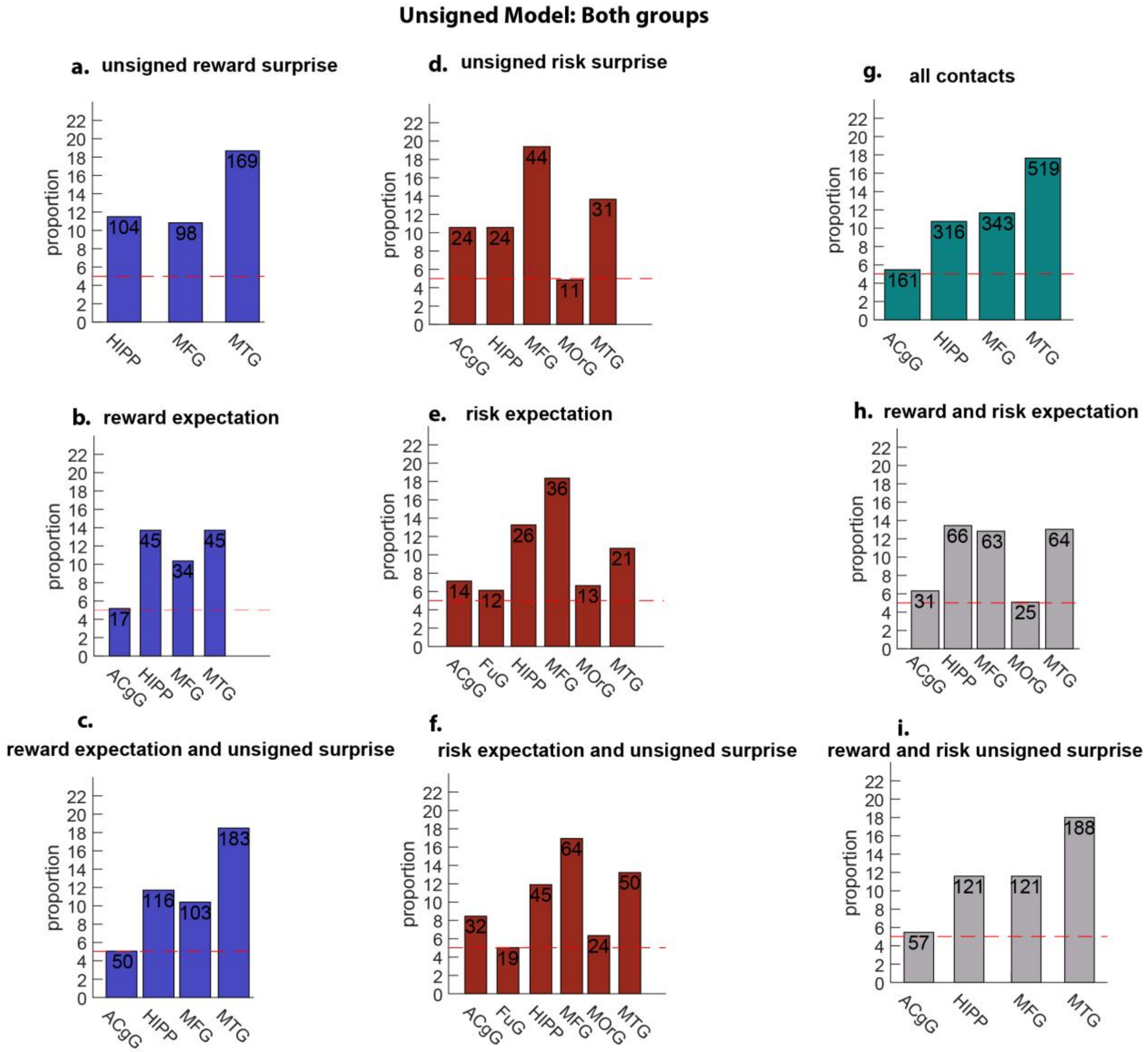


**Figure S7b. Unsigned Flat Brain: Proportion tests for Both Impulsivity Groups. (a)** Unsigned Reward Surprise. **(b)** Reward Value Expectation. **(c)** Reward Value Expectation and Unsigned Surprise. **(d)** Unsigned Risk Surprise. **(e)** Risk Value Expectation. **(f)** Risk Value Expectation and Unsigned Surprise. **(g)** All Contacts. **(h)** Reward and Risk Value Expectation. **(i)** Reward and Risk Unsigned Surprise.

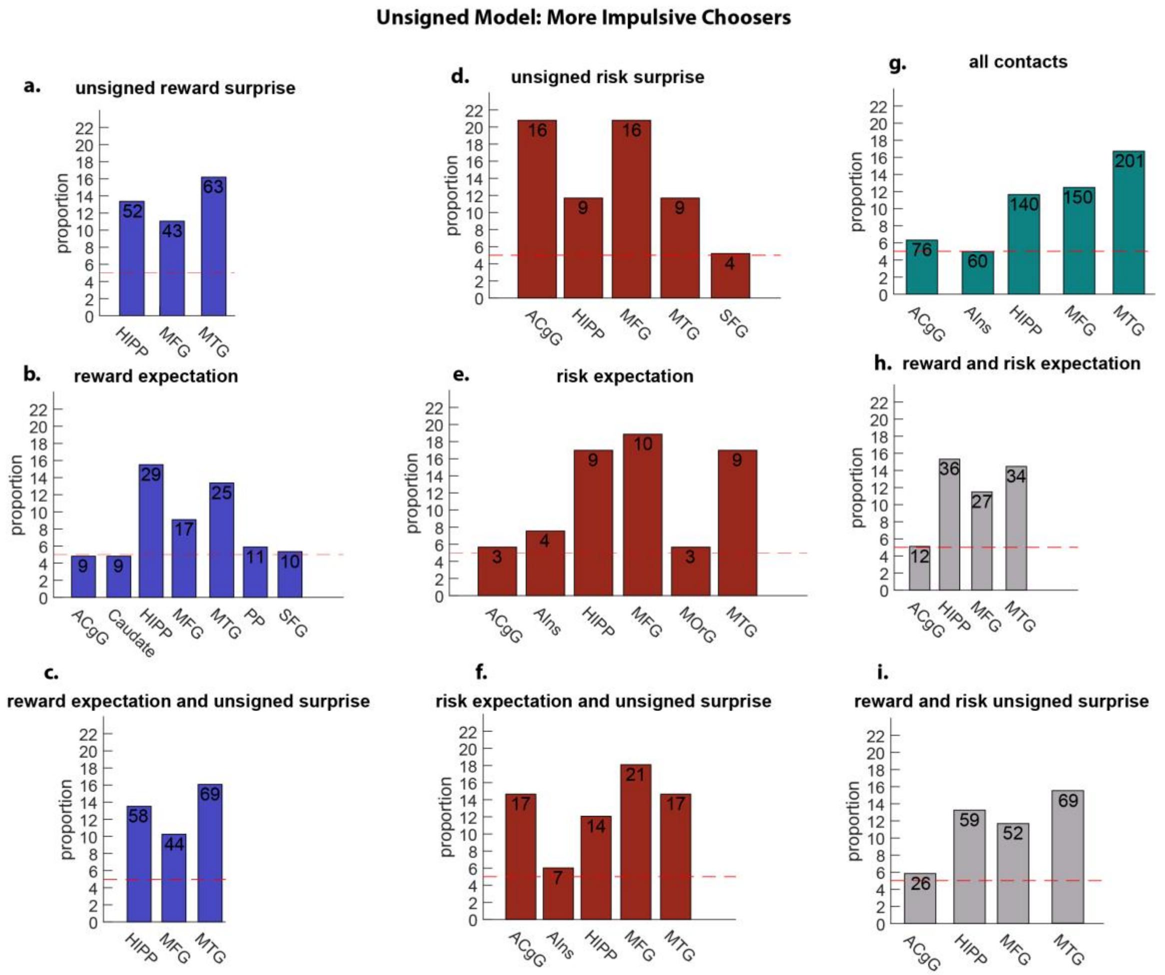


**Figure S7c. Unsigned Flat Brain: Proportion tests for More Impulsivity Groups. (a)** Unsigned Reward Surprise. **(b)** Reward Value Expectation. **(c)** Reward Value Expectation and Unsigned Surprise. **(d)** Unsigned Risk Surprise. **(e)** Risk Value Expectation. **(f)** Risk Value Expectation and Unsigned Surprise. **(g)** All Contacts. **(h)** Reward and Risk Value Expectation. **(i)** Reward and Risk Unsigned Surprise.

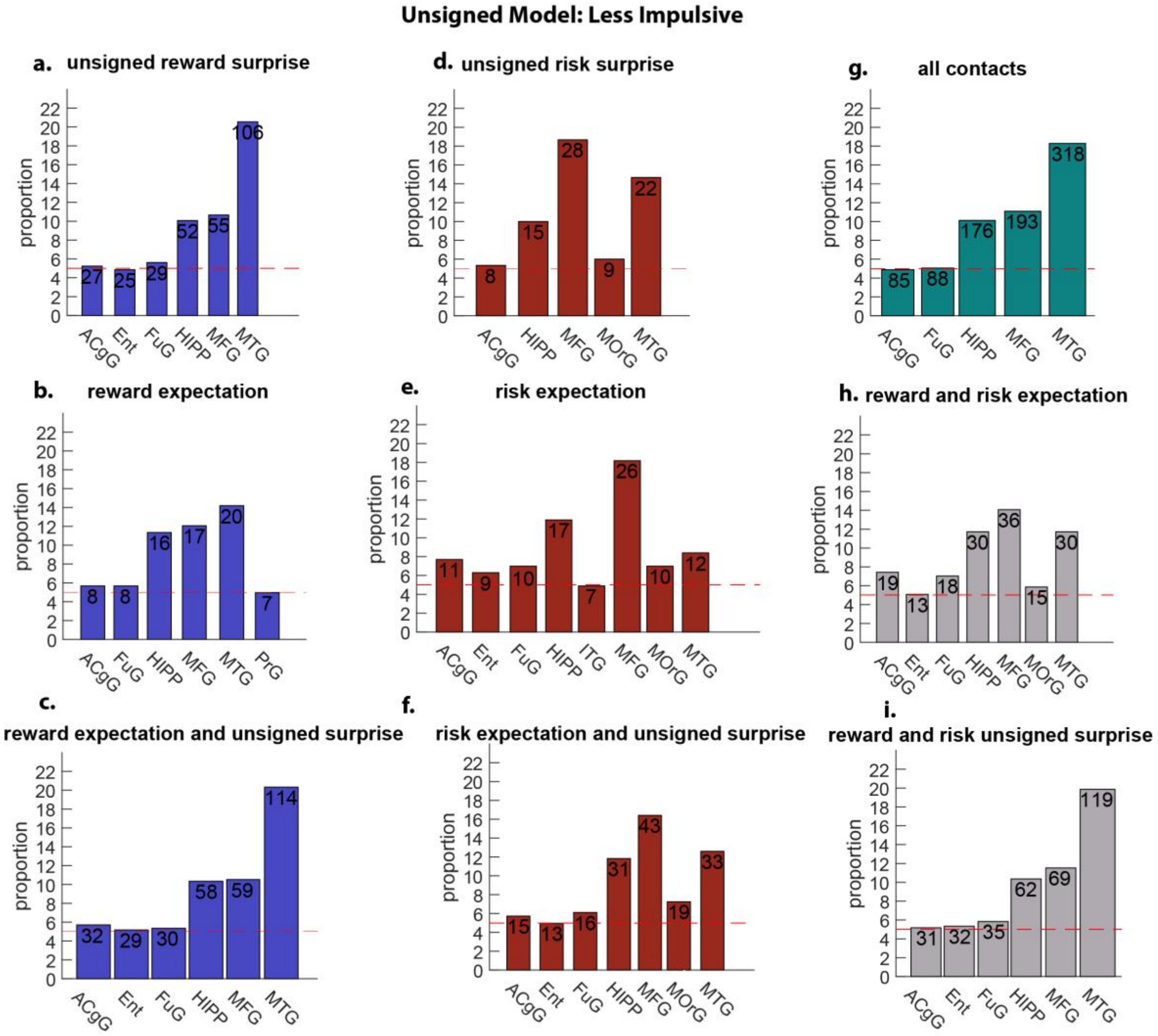


**Figure S7d. Unsigned Flat Brain: Proportion tests for Less Impulsivity Groups. (a)** Unsigned Reward Surprise. **(b)** Reward Value Expectation. **(c)** Reward Value Expectation and Unsigned Surprise. **(d)** Unsigned Risk Surprise. **(e)** Risk Value Expectation. **(f)** Risk Value Expectation and Unsigned Surprise. **(g)** All Contacts. **(h)** Reward and Risk Value Expectation. **(i)** Reward and Risk Unsigned Surprise.

**Table S3. Participant Information.** Patient Number, Gender, and Age.

| Patient | Gender | Age | Patient | Gender | Age | Patient | Gender | Age |
|---|---|---|---|---|---|---|---|---|
| 1 | M | 44 | 16 | M | 58 | 31 | F | 47 |
| 2 | F | 37 | 17 | F | 52 | 32 | M | 22 |
| 3 | F | 42 | 18 | M | 43 | 33 | M | 44 |
| 4 | F | 42 | 19 | F | 45 | 34 | M | 26 |
| 5 | F | 35 | 20 | M | 55 | 35 | M | 41 |
| 6 | M | 45 | 21 | F | 41 | 36 | F | 35 |
| 7 | M | 33 | 22 | M | 49 | 37 | F | 31 |
| 8 | M | 27 | 23 | M | 26 | 38 | F | 30 |
| 9 | M | 22 | 24 | M | 38 | 39 | F | 46 |
| 10 | M | 54 | 25 | M | 37 | 40 | F | 35 |
| 11 | M | 28 | 26 | F | 39 | 41 | F | 21 |
| 12 | M | 30 | 27 | F | 42 | 42 | F | 45 |
| 13 | M | 29 | 28 | M | 26 | 43 | F | 19 |
| 14 | M | 59 | 29 | F | 32 | | | |
| 15 | F | 50 | 30 | M | 35 | | | |

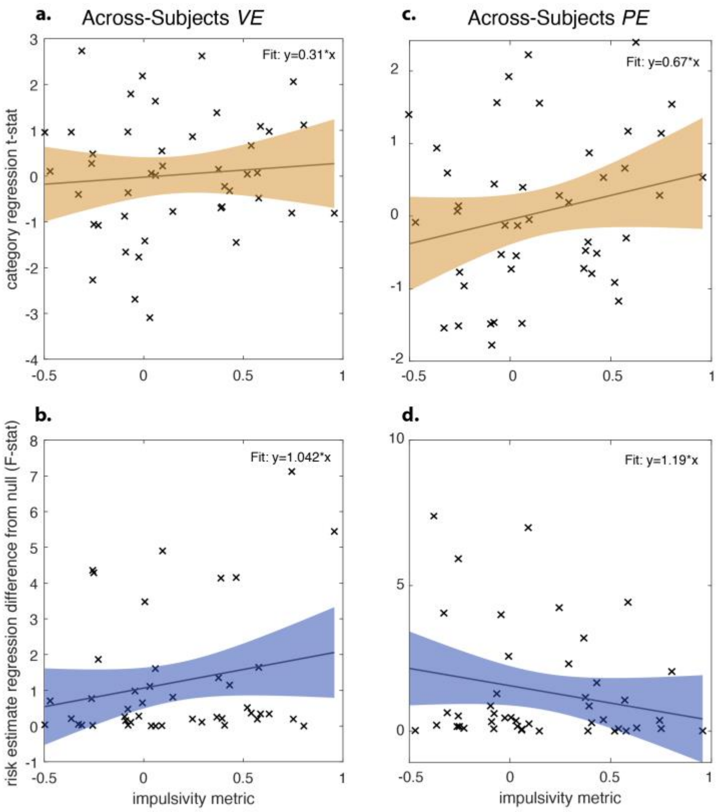


**Figure 8. Across-subjects results for Response Times, Value Expectation and Prediction Error analyses. (a)** t-statistic value for each subject for value expectation. **(b)** regression between risk estimate (F-stat) and impulsivity score (logKLD). **(c)** t-statistic value for each subject for prediction error. **(d)** regression between risk estimate (F-stat) and impulsivity score (logKLD).

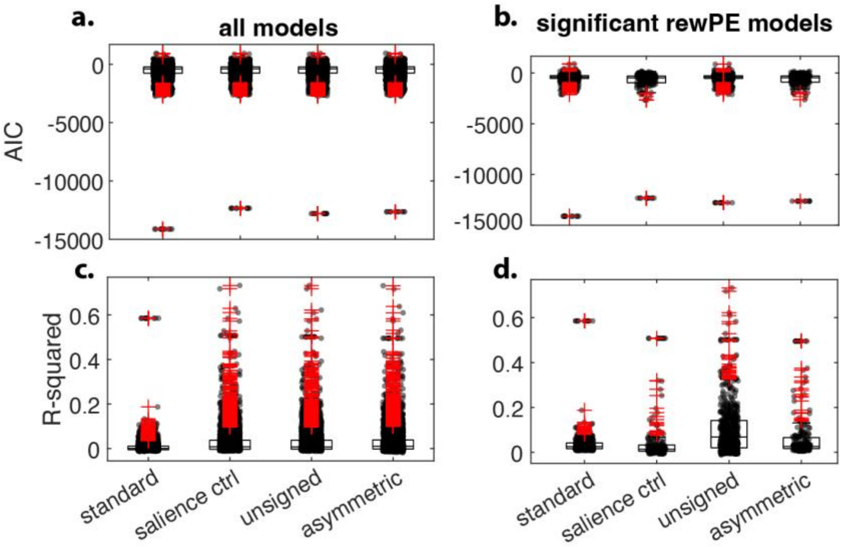


**Figure 9. All and Significant Reward PE models. (a)** AIC for the standard, salience control, unsigned, and asymmetric models: F(3) = 8.12, p = 0.000023. **(b)** R-Squared values for the standard, salience control, unsigned, and asymmetric models: F(3) = 15.48, p < 10-9.

**Table S4. Asymmetric Model.** Percentage and raw proportions of significantly encoded contacts for LI, MI, and both impulsivity groups. NA highlights non-significant regions.

| Encoding Regions<br>Less Impulsive Group | | | | | | | | | | | |
|---|---|---|---|---|---|---|---|---|---|---|---|
| **TD Variable** | **MTG** | **MFG** | **HIPP** | **FuG** | **ACgG** | **ENT** | **MOrG** | **AOrG** | **AMY** | **CO** | **Caudate** |
| **Risk PEs** | 27 (8%) | 17 (9%) | 19 (11%) | 13 (15%) | *NA* | *NA* | 19 (32%) | *NA* | *NA* | *NA* | *NA* |
| **RPEs** | 56 (18%) | 59 (31%) | 38 (22%) | *NA* | 18 (21%) | 21 (29%) | *NA* | *NA* | *NA* | *NA* | *NA* |
| **pRPEs** | 18 (6%) | 20 (10%) | 8 (5%) | *NA* | 8 (9%) | 7 (10%) | *NA* | 8 (32%) | 7 (21%) | *NA* | *NA* |
| **P Risk PEs** | 16 (5%) | *NA* | 9 (5%) | 7 (8%) | *NA* | 9 (12%) | 10 (17%) | *NA* | *NA* | *NA* | *NA* |
| **nRPEs** | 40 (13%) | 42 (22%) | 33 (19%) | *NA* | *NA* | 15 (21%) | *NA* | *NA* | *NA* | *NA* | *NA* |
| **N Risk PEs** | 13 (4%) | 13 (7%) | 13 (7%) | 6 (7%) | *NA* | *NA* | 13 (22%) | *NA* | *NA* | 8 (22%) | 7 (23%) |
| **Total Contacts** | **318** | **193** | **176** | **88** | **85** | **73** | **60** | **25** | **34** | **37** | **30** |

| Encoding Regions<br>More Impulsive Group | | | | | | | | | | | | | | |
|---|---|---|---|---|---|---|---|---|---|---|---|---|---|---|
| **TD Variable** | **MTG** | **MFG** | **HIPP** | **FuG** | **ACgG** | **ENT** | **MOrG** | **Ains** | **AMY** | **PP** | **OrIFG** | **POrG** | **TrIFG** | **STG** |
| **Risk PEs** | 17 (8%) | 22 (15%) | 8 (6%) | *NA* | 19 (25%) | *NA* | *NA* | 9 (15%) | *NA* | *NA* | *NA* | *NA* | *NA* | *NA* |
| **RPEs** | 73 (36%) | 43 (29%) | 59 (42%) | *NA* | *NA* | *NA* | 21 (47%) | *NA* | *NA* | *NA* | *NA* | *NA* | *NA* | *NA* |
| **pRPEs** | 18 (9%) | 8 (5%) | 17 (12%) | *NA* | *NA* | *NA* | *NA* | *NA* | 10 (37%) | 8 (23%) | *NA* | *NA* | *NA* | *NA* |
| **P Risk PEs** | *NA* | 10 (7%) | *NA* | *NA* | 12 (16%) | 4 (12%) | *NA* | 6 (10%) | *NA* | *NA* | 4 (24%) | 4 (14%) | 4 (13%) | *NA* |
| **nRPEs** | 59 (29%) | 35 (23%) | 53 (38%) | *NA* | 15 (20%) | *NA* | 19 (42%) | *NA* | *NA* | *NA* | *NA* | *NA* | *NA* | *NA* |
| **N Risk PEs** | 15 (7%) | 13 (9%) | 5 (4%) | *NA* | 8 (11%) | *NA* | *NA* | 6 (10%) | *NA* | 5 (14%) | *NA* | 5 (18%) | *NA* | 5 (21%) |
| **Total Contacts** | **201** | **150** | **140** | **26** | **76** | **34** | **45** | **60** | **27** | **35** | **17** | **28** | **32** | **24** |

| Encoding Regions<br>Both Impulsive Group | | | | | | | | | |
|---|---|---|---|---|---|---|---|---|---|
| **TD Variable** | **MTG** | **MFG** | **HIPP** | **FuG** | **ACgG** | **ENT** | **MOrG** | **PORG** | **AMY** |
| **Risk PEs** | 44 (8%) | 39 (11%) | 27 (9%) | *NA* | 27 (17%) | *NA* | 25 (24%) | *NA* | *NA* |
| **RPEs** | 129 (25%) | 102 (30%) | 97 (31%) | *NA* | *NA* | *NA* | *NA* | *NA* | *NA* |
| **pRPEs** | 36 (7%) | 28 (8%) | 25 (8%) | *NA* | *NA* | *NA* | *NA* | *NA* | 17 (28%) |
| **P Risk PEs** | 19 (10%) | 14 (4%) | 12 (4%) | 9 (%) | 16 (10%) | 13 (12%) | 13 (12%) | *NA* | *NA* |
| **nRPEs** | 99 (19%) | 77 (22%) | 86 (22%) | *NA* | *NA* | *NA* | 32 (30%) | *NA* | *NA* |
| **N Risk PEs** | 28 (5%) | 26 (8%) | 18 (8%) | *NA* | 12 (7%) | *NA* | 16 (15%) | 11 (21%) | *NA* |
| **Total Contacts** | **519** | **343** | **316** | **114** | **161** | **107** | **105** | **52** | **61** |

**Table S5. Unsigned Model.** Percentage and raw proportions of significantly encoded contacts for LI, MI, and both impulsivity groups. NA highlights non-significant regions.

| | Encoding Regions Less Impulsive Group | | | | | | | Encoding Regions More Impulsive Group | | | | | Encoding Regions Both Impulsive Group | | | |
|---|---|---|---|---|---|---|---|---|---|---|---|---|---|---|---|---|
| **TD Variable** | **MTG** | **MFG** | **HIPP** | **FuG** | **ACgG** | **ENT** | **MOrG** | **MTG** | **MFG** | **HIPP** | **ACgG** | **SFG** | **MTG** | **MFG** | **HIPP** | **ACgG** |
| **Unsigned RPE** | 106 (33%) | 55 (28%) | 52 (30%) | 129 (33%) | 27 (32%) | *NA* | *NA* | 63 (31%) | 43 (29%) | 52 (37%) | *NA* | *NA* | 169 (33%) | 98 (29%) | 104 (33%) | *NA* |
| **Unsigned Risk PE** | 22 (7%) | 28 (15%) | 15 (9%) | *NA* | 8 (9%) | NA | 9 (15%) | 9 (4%) | 16 (11%) | 9 (6%) | 16 (21%) | 4 (11%) | 31 (6%) | 44 (13%) | 24 (8%) | 24 (15%) |
| **Unsigned Risk PE & RPE** | 119 (37%) | 69 (36%) | 62 (35%) | 35 (40%) | 31 (36%) | 32 (44%) | *NA* | 69 (34%) | 52 (35%) | 59 (42%) | 26 (34%) | *NA* | 188 (36%) | 121 (35%) | 121 (38%) | 57 (35%) |
| **Total Contacts** | **318** | **193** | **176** | **88** | **85** | **73** | **60** | **201** | **150** | **140** | **76** | **24** | **519** | **343** | **316** | **161** |

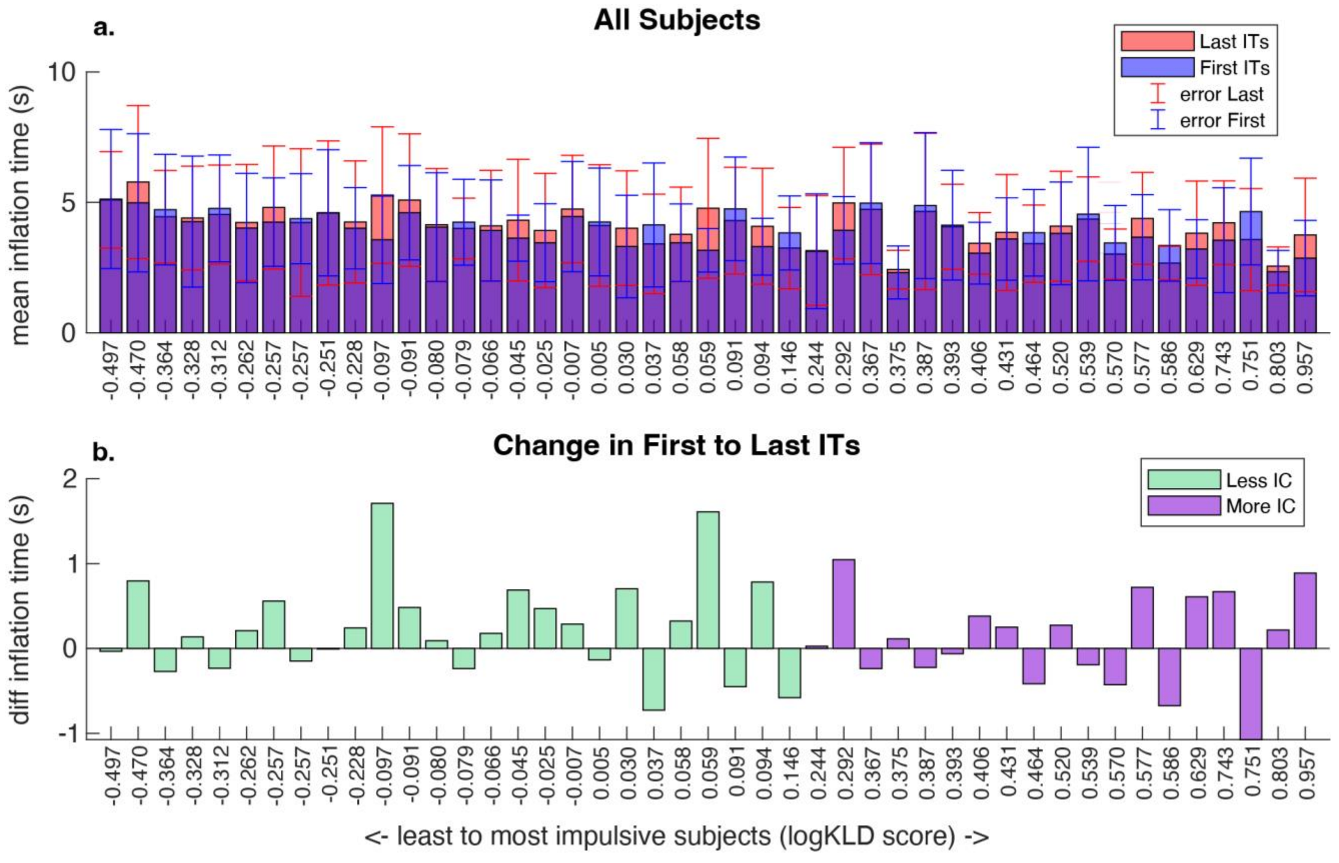


**Figure 10. First and Last 50 Trial Inflation Times. (a)** Histogram of first 50 (blue) and last 50 (red) trials from BART ordered by impulsivity. Standard deviation errors bars for both sets of trials. **(b)** Difference in inflation time from first to last 50 trials. Positive inflation time difference indicates an increase in inflation time as the task goes on. Negative inflation time difference indicates a decrease in inflation time as the task goes on. Ordered from least to most impulsive choosers.

### Original TD model encoding:

**Tests for overall encoding of each type of TD model (reward vs risk):**
Proportion of contacts significantly encoding reward vs risk variables: (79.48%, 23.45%) ($\chi^2$(1) = 1014.42, p = 0.00000000).
Proportion of contacts significantly encoding Reward VE vs Reward PE: (21.54%, 47.73%) ($\chi^2$(1) = 240.73, p = 0.00000000).
Proportion of contacts significantly encoding Risk VE vs Risk PE: (14.13%, 11.81%) ($\chi^2$(1) = 3.89, p = 0.0487).

**Tests for REWARD model encoding by impulsivity classification:**
Proportion of contacts significantly encoding value expectation (more impulsive, less impulsive): (13.72%, 6.44%) ($\chi^2$(1) = 48.82, p = 0.0000000).
Proportion of contacts significantly encoding surprise (PE) (more impulsive, less impulsive): (26.02%, 26.30%) ($\chi^2$(1) = 0.03, p = 0.855).
Proportion of contacts significantly encoding value and surprise (PE) (more impulsive, less impulsive): (10.46%, 4.14%) ($\chi^2$(1) = 49.56, p = 0.00000000).

**Tests for RISK model encoding by impulsivity classification:**
Proportion of contacts significantly encoding risk value expectation (more impulsive, less impulsive): (4.35%, 8.06%) ($\chi^2$(1) = 18.6, p = 0.000016).
Proportion of contacts significantly encoding risk surprise (PE) (more impulsive, less impulsive): (5.57%, 7.22%) ($\chi^2$(1) = 3.62, p = 0.0571).
Proportion of contacts significantly encoding risk value and surprise (PE) (more impulsive, less impulsive): (1.49%, 1.23%) ($\chi^2$(1) = 0.42, p = 0.517).

**Tests for REWARD AND RISK model encoding by impulsivity classification:**
Proportion of contacts significantly encoding reward and risk value expectation (more impulsive, less impulsive): (1.09%, 1.01%) ($\chi^2$(1) = 0.05, p = 0.824).
Proportion of contacts significantly encoding reward and risk surprise (PE) (more impulsive, less impulsive): (1.77%, 2.69%) ($\chi^2$(1) = 3.08, p = 0.0794).
Proportion of contacts significantly encoding reward and risk value and surprise (PE) (more impulsive, less impulsive): (0.68%, 0.11%) ($\chi^2$(1) = 7.08, p = 0.0.778).

**Tests for hemispheric differences:**
Proportion of contacts significantly encoding reward model variables in the left hemisphere (left hemisphere, right hemisphere): (44.02%, 29.21%), ($\chi^2$(1) = 76.94, p = 0.000000).
Proportion of contacts significantly encoding risk model variables in the left hemisphere (left hemisphere, right hemisphere): (14.47%, 11.53%), ($\chi^2$(1) = 6.24, p = 0.0125).
Proportion of contacts significantly encoding reward model variables in the left hemisphere (left hemisphere, right hemisphere) in more impulsive subjects: (28.46%, 9.29%), ($\chi^2$(1) = 201.51, p = 0.000000).
Proportion of contacts significantly encoding risk model variables in the left hemisphere (left hemisphere, right hemisphere) in more impulsive subjects: (7.20%, 2.24%), ($\chi^2$(1) = 46.45, p = 0.000000).
Proportion of contacts significantly encoding reward model variables in the left hemisphere (left hemisphere, right hemisphere) in less impulsive subjects: (15.56%, 19.92%), ($\chi^2$(1) =10.44, p = 0.00123).
Proportion of contacts significantly encoding risk model variables in the left hemisphere (left hemisphere, right hemisphere) in less impulsive subjects: (7.27%, 9.29%), ($\chi^2$(1) = 4.29, p = 0.0383).
